# Compositional and interpretable representation of histology using AI foundation models and sparse autoencoders

**DOI:** 10.64898/2026.06.03.725182

**Authors:** Ziyuan Zhao, Zoltan Maliga, Emmanuel C. Ogbonna, Soheil R. Talemi, Shannon Coy, Andréanne Gagné, Kapongo Lumamba, Isaac H. Solomon, Sandro Santagata, Adrie J.C. Steyn, Threnesan Naidoo, Peter K. Sorger

## Abstract

Light microscopy of tissue sections stained with hematoxylin and eosin (H&E) has been the foundation of histopathology for over 150 years and remains essential for diagnosis and research. The development of high-plex spatial profiling approaches able to measure protein and RNA expression at single-cell resolution augments but does not replace H&E imaging, even in research. Computational pathology (CPath) models based on deep learning promise to further increase the value of H&E imaging but interpreting these models in biological terms remains challenging. As a result, they are not widely used in spatial profiling studies. Here we describe a human-in-the-loop computational framework that leverages CPath foundation models (FMs) and sparse autoencoders (SAEs) to decompose FM embeddings and automatically identify diverse, human-interpretable histopathology features in H&E images. When FM-SAE modeling was applied to pulmonary diseases such as tuberculosis and lung cancer, human-machine interaction augmented and accelerated expert interpretation. Moreover, the resulting annotations provide a morphology-aware approach to integrating 2D and 3D mesoscale tissue architectures with molecular spatial profiling.

## INTRODUCTION

Tissues are organized at multiple spatial scales from intracellular organelles and macromolecular complexes to functional tissue units (FTUs)^1^ measuring millimeters to centimeters in length and comprising multiple cell types and acellular structures. For over 150 years, histopathologists have studied the organization, morphology, and functions of FTUs as well as their association with disease progression and mortality^2^, primarily via examination of tissues stained with dyes such as hematoxylin and eosin (H&E) or with antibodies via immunohistochemistry (IHC)^3^. Early efforts to replicate this process computationally by using defined sets of morphometric features (e.g. nuclear density or ellipticity) date back to the 1960s^4^, but it is only in the past decade that the potential of computational pathology (CPath)^5^ has become apparent, thanks to rapid advances in AI/ML models^6,7^. This has yielded deep learning (DL) models for identification of specific structures in H&E images (e.g., glands, immune aggregates, and vasculature) using supervised^8–11^ or weakly supervised learning approaches^12–14^. These approaches commonly involve manual annotation of the training data, which requires substantial expertise and effort, limiting such approaches to specific structures and diseases. These limitations have motivated the development of unsupervised^15–17^ approaches in which task-driven annotation of images is replaced by data-driven featurization of whole-tissue composition for downstream computational tasks^18^ or human review^19,20^.

More recently, foundation models (FMs) trained in an unsupervised manner on very large sets of digital H&E images have been developed as a general backbone for CPath^21^. FMs can encode general representations of tissue morphology for a wide range of tasks, but current benchmarking and deployment efforts focus primarily on diagnosis performed at the level of WSIs, such as tumor classification, staging, and survival prediction^22–25^, rather than systematic morphological analysis for biological discovery. With an FM such as UNI^26^, tissue morphology is typically represented by embedding tiled image patches using a vision transformer and then interpreting the embeddings by applying discrete clustering or linear classifiers. Standard workflows enforce a single, exclusive label on each image patch that is fundamentally at odds with the variable appearance and overlap of histomorphologies. The field has recognized this complexity, leading to the application of alternative representations of tissues, with diffusion, pseudotime, and latent factor modeling^27,28^, or mixtures of cellular motifs or inferred dynamics^29–32^.

In parallel with the development of CPath models of H&E images, the past decade has witnessed the development of a wide variety of highly multiplexed spatial profiling technologies able to profile tens to thousands of proteins, mRNAs, and metabolites in intact tissue at cellular to subcellular resolution^33,34^. It is often assumed that these data can be used directly to infer biologically relevant tissue features (e.g., by trajectory inference)^35^ with minimal input from histopathology. However, this misses an opportunity to leverage foundational knowledge of pathology and pathophysiology derived from 150 years of inspection of H&E images^36^. Histopathologists are highly trained to identify tissue features and abnormalities at both gross and microscopic scales, including architectural (multicellular organization), cytological (cell-level morphology) and subcellular levels (the shapes and distributions of organelles).

This information is used to formulate a diagnosis based on established clinicopathologic associations between tissue features and disease states. The application of CPath to H&E images acquired in parallel with high-plex spatial profiles^37^ represents a powerful opportunity to better create a bridge between molecular measurements, functional properties of tissues, and pathologic knowledge of disease. This is particularly important when acellular structures such as basement membranes, elastic fibers, and collagen etc. play a critical role: these tissue constituents are visible in H&E images but not in transcriptional profiles.

There are few studies to date linking high-plex spatial profiling, CPath, and foundational knowledge of histopathology that has been summarized by international classification^38^ and staging^39^ guidelines. In part this is because FM CPath outputs are not immediately human interpretable. It is also the case that diagnosis and staging – a primary focus of current FM models – is focused on histological features useful in differential diagnosis, prognostication, or treatment response prediction. This omits many features of potentially great scientific value^40,41^. In a research setting, these features represent a rich basis for discovery of new biological phenomena, novel disease entities, and the refinement of diagnostic criteria^42^ and it is frequently necessary to annotate research specimens at a finer level of detail than for clinical diagnosis.

In this paper we tackle these issues using a human-in-the-loop framework that combines pretrained histopathology FMs with sparse autoencoders (SAEs). We show that this approach makes it possible to align FM model embeddings with histopathology knowledge as interpreted by multiple board-certified pathologists. This enabled labeling >90% of the area of whole-slide H&E images with biologically relevant features and interpretation of highly multiplexed tissue images (CyCIF)^43^ from adjacent tissue sections. We focus on two lung diseases, tuberculosis (TB) and lung cancer, that exhibit common features of inflammation, immune aggregation, and tissue remodeling. Le et al. (2025)^44^ also used SAEs to decompose FM embeddings, but with the primary goal of better understanding how models function. In contrast, our focus is on model interpretability and utility in and out of domain as a means of quantifying the variable structure of human tissues at scale.

## RESULTS

### Clustering of foundation model embeddings fails to capture compositional histomorphology

To explore the use of a FM in automated annotation of complex histomorphologies, we processed multiple datasets comprising H&E images of conventional 4-5 µm tissue sections prepared from formalin-fixed paraffin-embedded (FFPE) specimens recovered by surgery or autopsy; H&E staining and digitization were performed using standard histopathology protocols at multiple medical centers (**Supplementary Table 1**). Each dataset was designed to represent an increasingly diverse set of images. Dataset 1 (DS1) comprised 40 sequential H&E WSIs spaced every 15 µm, originating from a single tissue block selected from the lung resection of an untreated TB patient; each H&E image had a corresponding adjacent 39-plex CyCIF image^45^. DS2 comprised 15 WSIs from sections of 10 different tissue blocks from the same patient; DS3 comprised 20 WSIs collected at autopsy or pneumonectomy of advanced and treated TB patients. In addition, we processed images from the literature that included 12 sections from resections of lung adenocarcinomas (DS4)^46^ and 5 sections of sarcoidosis (DS5)^47^, a non-infectious lung granulomatous disease of unknown etiology.

Inspection of DS1 by multiple board-certified pathologists, including a thoracic pathologist and an expert in TB pathology, from three institutions (collectively subject matter experts; SMEs) confirmed the presence of a wide range of histomorphologies, including disease-relevant features such as granulomas, lymphoid aggregates (LA) and tertiary lymphoid structures (TLS), as well as normal lung parenchymal structures such as blood vessels, airways, and alveoli (**Fig. 1a**). Granulomas are the hallmark lesion of pulmonary TB and consist of organized aggregates of macrophages surrounded by a rim of lymphocytes, typically with a central region of caseous necrosis (tissue death with a crumbly “cheese-like” appearance) and often bordered by fibrosis in more developed lesions^48,49^. To analyze these and other features in a data-driven manner using an FM, we tiled overlapping ∼120 x 120 μm (448 x 448 pixel) patches from WSIs derived from DS1 to generate a set of ∼2 × 10^6^ image patches that were then encoded using UNI. UNI is a histopathology FM^26^ trained by self-supervised learning on >10^5^ H&E WSIs representing a wide range of diseases and tissue types to generate 1024-dimensional embeddings. As previously shown^23^, UNI embeddings provide a compact representation of tissue morphology at the spatial scale of a patch or smaller.

**Figure 1.**
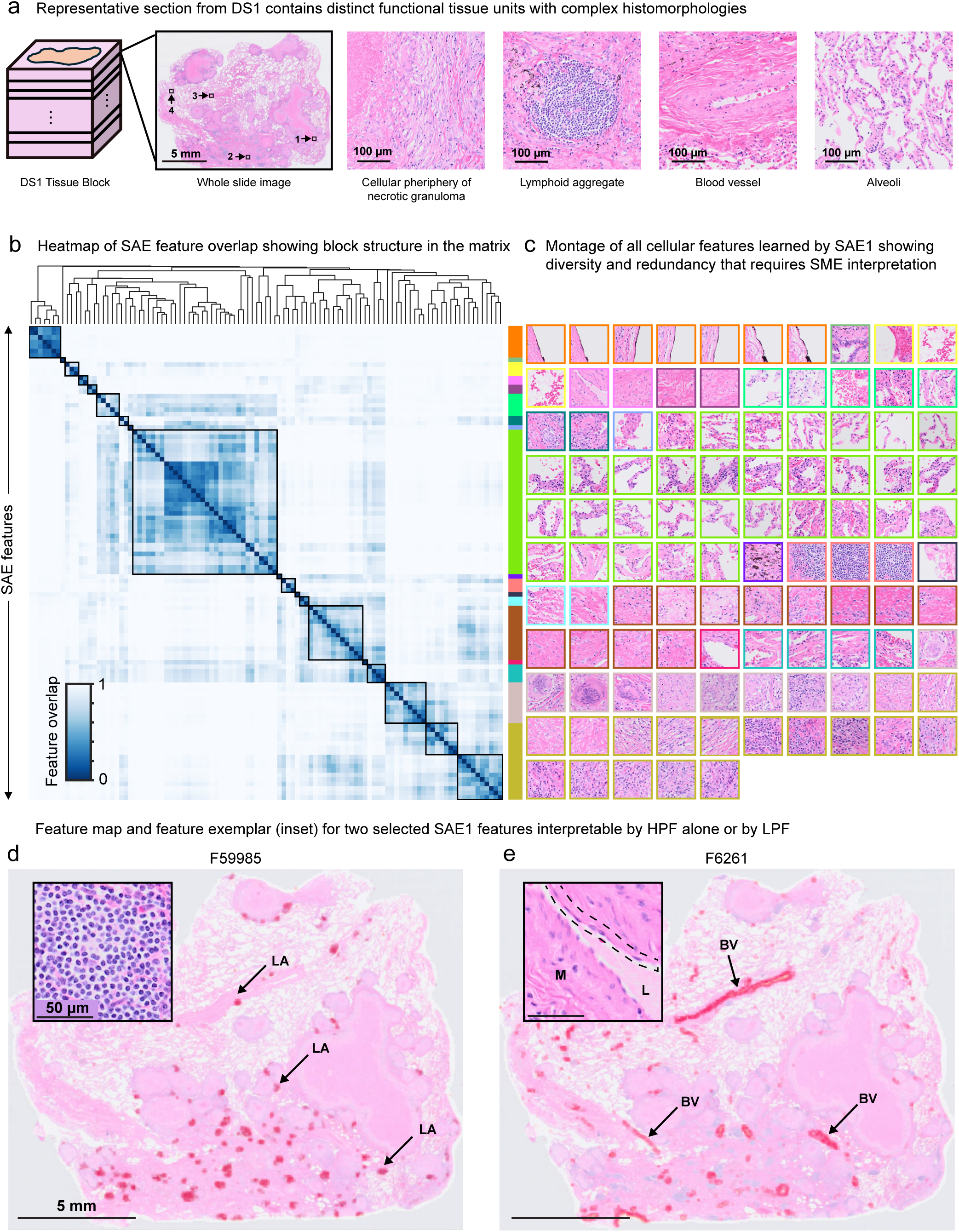
Defining complex histomorphologies in a 3D H&E dataset of human pulmonary tuberculosis using FM-SAE. **(a)** Cartoon of serial sectioning plan for FFPE tissue block to collect DS1 training data. Representative H&E WSI containing four FOVs (indicated). Inset, each FOV illustrates distinct histomorphologies as indicated. **(b)** Heatmap, spatial overlap for all 361 cellular features sorted by hierarchical clustering (method). Morphologic clusters are outlined on the heatmap and coded (colormap, right) to match (c). **(c)** H&E image montage, highest scoring patches for each sorted SAE1 feature and outlined as indicated in the colorbar from (b). Representative feature maps, H&E WSI with embedded image patch feature score (red) and, inset, exemplar image patch from (c). **(d)** Feature F59985, lymphocytic aggregate, as indicated (arrows, LA). **(e)** Feature 6261, large blood vessel (BV), as indicated (arrows, BV) in WSI. Inset, labeling BV features: lumen (L), muscularis (M), and endothelial cells (dashed line).

Clustering is the standard approach to organizing FM embeddings. We compared two widely used methods: K-means^50^ clustering and graph-based Leiden community detection^51^. In the former approach, we set K = 25 (**Supplementary Fig. 1a**) as a trade-off between the morphological diversity of lung tissue and the practical constraints of pathologist review, providing recognizable tissue architectural features when projected onto WSIs (**Supplementary Fig. 1b**). Default parameters for Leiden community detection^51^ generated 21 clusters with similar overall UMAP (**Supplementary Fig. 1c**) and WSI spatial profiles (**Supplementary Fig. 1d**). This level of cluster granularity is consistent with existing applications of histopathology FMs^18,52^.

SMEs then attempted to interpret the histomorphologic feature represented in each cluster by reviewing a standardized feature report. The report included 20 representative image patches (what a pathologist might call a “high-power field” or “high magnification”)^53^ and patches mapped onto 10 WSIs (“low-power fields” or “low magnification”) to provide tissue context (**Supplementary Fig. 1e**). We found the clusters fell into three general classes: (1) tissue regions rich in cells of different types (19 K-means clusters); (2) regions with few intact cells that stained a uniform pink or purple and corresponded to granuloma cores (3 clusters); (3) empty space with occasional debris corresponding to tissue edges and alveoli with associated debris (3 clusters; **Supplementary Table 2**; **Supplementary Fig. 1f**). Focusing on cell-rich features^49,54^, SMEs found that a single image patch commonly contained multiple histomorphologies including varying cell types and acellular structures in different architectural arrangements; this has previously been described for images of mesothelioma processed using a similar DL model^15^ and is illustrated in **Supplementary Fig. 1g** for representative patches from lymphoid aggregate (K-means cluster C17) and fibrosis (C7). C17 patches contained round cells that stain dark purple with H&E, which are lymphoid cells (primarily B and T cells) matching the patch label. However, other C17 patches contain varying levels of interleaving acellular pink fibrosis, and yet others had interleaved domains of lymphocytes and fibrosis. The converse was true of C7: fibrosis dominated the patches, but many also contained substantial numbers of lymphoid cells. This type of intermixing was observed for a wide variety of cell types, ECM components, and multicellular structures (e.g., blood vessels, granulomas), resulting in ambiguous cluster labels. We hypothesized that the difficulty in assigning unambiguous labels to clusters of FM embeddings arose not from a limitation of FMs themselves, but from the requirement in conventional discrete clustering that any image patch, even one composed of multiple histomorphologies, be assigned to a single cluster. This 1:1 mapping, we postulated, misrepresents the underlying reality that histologic regions at the size of image patches (ca. 100-400 cells) commonly contain mixtures of multiple cell types and tissue structures.

### Sparse autoencoders decompose foundation model embeddings into interpretable histomorphologic features

To overcome the limitations of discrete clustering, we applied a compositional approach to the representation of FM embeddings. This could be implemented, in principle, using any latent factor modeling method that allows a single patch to be represented computationally as a combination of multiple elements. We chose a sparse autoencoder (SAE)^44,55^ that learns to decompose patch embeddings into weighted sums of features, with non-negative weights referred to as “feature scores” (**Supplementary Fig. 2a**). When an SAE with a single-layer neural network was trained on UNI embeddings from DS1, 361 highly active features were obtained after filtering (see methods and **Supplementary Fig. 3a** for filtering details). To facilitate visual review, these features were organized using hierarchical clustering based on pairwise *feature overlap* (**Supplementary Fig. 2b**). We defined feature overlap using an Intersection over Union (IoU) metric, calculated as the number of patches with high scores for both features divided by the number of patches with a high score for either feature.

Because patches were densely sampled from a regular grid, each region of tissue was represented by multiple overlapping patches, and the set-level IoU also corresponded in many cases to spatial co-localization. Features were then ordered by hierarchically clustering the feature-overlap matrix into initial clusters (**Fig. 1b, Supplementary Fig. 3b**). The top-scoring patch (“exemplars”) from each feature was used to generate a montage of all features (**Fig. 1c, Supplementary Fig. 3c**), and feature reports were generated to facilitate SME review of top-scoring image patches and the spatial distribution of feature scores in WSIs (**Supplementary Fig. 3d**).

SMEs judged some FM-SAE features to be readily interpretable based on examination of top-scoring patches and WSI context. For example, feature F59985 corresponded to aggregates of lymphoid cells, many of which were localized to clusters around granulomas (**Fig. 1d**). Similarly, high-magnification image patches and tissue distribution of feature F6261 were consistent with larger-caliber blood vessels (**Fig. 1e**). However, it remained challenging to interpret patches with mixed histomorphologies based on patch-level feature scores alone. To address this, H&E images were overlaid with (partially transparent) maps depicting the saliency of each feature at the level of tokens, which are approximately the size of a single cell. This was accomplished by leveraging the vision transformer architecture used by the UNI FM, which encodes 32 x 32 pixel (8.8 x 8.8 µm) regions of a patch as individual tokens prior to aggregating all 196 tokens into patch-level embeddings. Each patch could in principle contain up to 196 different types of tokens, but within high-scoring patches for all SAE features examined, we found that it was sufficient to consider only two to three types of tokens, with the third often corresponding to “all other features.” To identify these token-level features, we performed K-means clustering of token embeddings with K = 3 (this parameter can be adjusted depending on the level of histological intermixing). For any patch scored by a feature, we generated a saliency map by applying its corresponding K-means classifier and mapping cluster assignments onto the patch (**Supplementary Fig. 2c**).

Saliency maps illustrate the composability of FM-SAE features. For example, when we selected image patches containing a range of mixed morphologies, with scores ranging from 0 to 0.89 for feature F27188 (lymphoid aggregate) and from 0 to 0.84 for F28301 (fibrosis), patch-level feature scores reflected the fraction of tokens associated with each feature (**Fig. 2b).** Leveraging this observation, we iteratively reviewed token-level saliency maps to merge token clusters into 18 groups and generate a dictionary of token labels (**Supplementary Fig. 4**). Saliency maps and token-level labels were particularly informative when a feature corresponded to only part of a patch (e.g., respiratory epithelium vs. “other”; **Fig. 2c)** or when multiple histomorphologies were associated with a feature. For example, top-scoring patches for FM-SAE feature F959, labeled by SMEs “inflamed tissue,” comprised varying mixtures of reactive pneumocytes, lymphocytes, and fibrosis (**Fig. 2d**). Pneumocytes are specialized epithelial cells involved in gas exchange lining the walls of alveoli that exhibit reactive changes in the context of inflammation and tissue repair, consistent with the designation “inflamed.”

**Figure 2.**
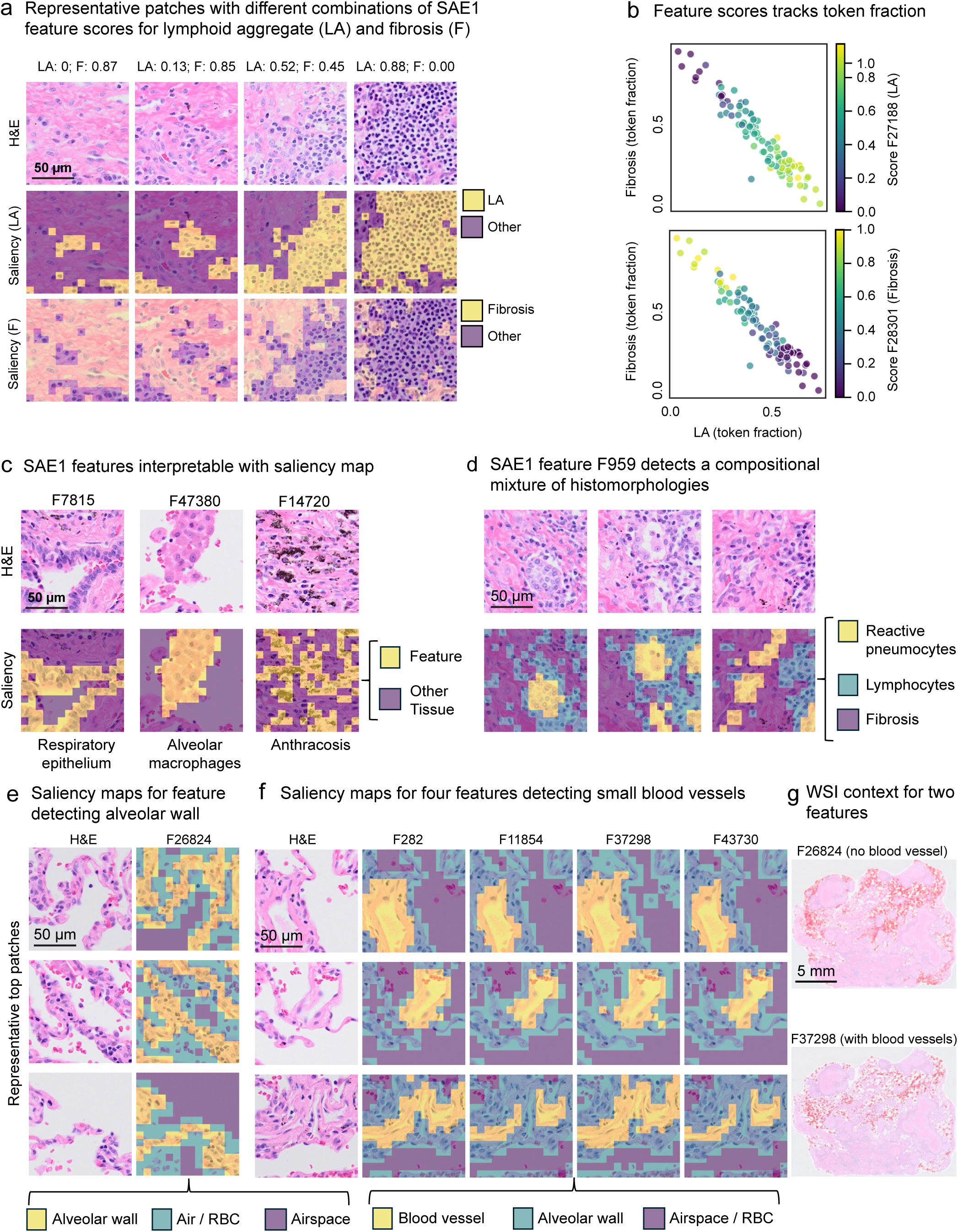
SAE detects composable and interpretable histomorphologies with feature-specific saliency maps. **(a)** FOVs with range of feature scores for F28301 (F: lung fibrosis) and F59985 (LA: lymphoid aggregate) (LA). Top, H&E image alone. Middle, H&E with image tokens shaded for LA (yellow) or other (violet) saliency. Bottom, H&E with image tokens shaded for fibrosis (yellow) or other (violet) saliency. **(b)** Scatterplot, fraction of image tokens positive for F28301 or F59985 in representative image patches (n=100) containing both. Color indicates normalized feature score for F28301 (top) or F27188 (bottom). **(c)** Top, exemplar H&E images for F7815 (respiratory epithelium), F47380 (intra-alveolar macrophages), and F14720 (anthracosis). Bottom, H&E with image tokens shaded for target (yellow) or other (violet) saliency. **(d)** Top, representative H&E images for F959, and bottom, image tokens shaded for feature-specific token clusters: reactive pneumocyte (yellow), lymphocyte (blue), and fibrosis (violet) saliencies. **(e)** Left, representative H&E images for F26824 (alveolar epithelium), right, image tokens shaded for feature-specific token clusters: alveolar wall (yellow), airspace with RBCs (blue) and airspace (violet). **(f)** Left. Exemplar H&E images with high scores for F282, F11854, F37298, and F43730. Right, token shading with feature-specific token clusters for blood vessel (yellow), alveolar wall (blue), and airspace (violet). **(g)** Representative H&E WSI with image patches colored red for, top, F26824 (alveolar parenchyma) and, bottom, F37298 (alveolar parenchyma with blood vessels).

In a second example, exemplar patches for F26824, labeled “alveolar wall” (comprising pneumocytes, associated capillaries, and interstitial tissue as well as airspace (**Fig. 2e**), differed visually from exemplar patches for features F282, F11854, F37298, and F43730, which also mapped to lung parenchyma. Saliency maps showed that the latter four FM-SAE features contained tokens corresponding to blood vessels, alveolar wall, and airspace (**Fig. 2f**). Thus, the key distinction between F26824 and the other four parenchymal features was the presence or absence in the patch of a small blood vessel (an arteriole or venule) in cross section. Because lung parenchyma contains extensive and regularly spaced vasculature, subdividing the tissue into patches will necessarily result in some patches containing arteriole or venules in cross-section and others not. All five parenchyma features therefore annotate different aspects of the same tissue and are spatially intermixed in WSIs (**Fig. 2g**). More generally, token-level saliency maps clarified relationships among FM-SAE features containing related but non-identical cell and tissue components. They also illustrated the multiple spatial scales at which histological labels are applied, including single cells (e.g. reactive pneumocytes), multicellular tissue features (e.g., alveolar wall), and FTUs (e.g., alveoli or blood vessels).

The key insight from this analysis was that assigning multiple feature scores to each image patch and visualizing the spatial distribution of these scores using token-level saliency maps helped align FM model outputs to existing knowledge about lung architecture. This knowledge was familiar to the SME team, but FM-SAE modeling made it more accessible to the computational team tasked with annotating large numbers of H&E images. The model also made the task of subdividing tissue into individual biological elements substantially faster and less laborious (see discussion).

### Compositional histomorphologic features map tissue architecture across whole-slide images

Overall, 356 (99%) FM-SAE features could be associated with established histopathology labels (**Supplementary Table 3**), and these features could be further consolidated with input from SMEs into 28 broad categories (**Fig. 3a, Supplementary Fig. 4, Supplementary Table 4**). These labels were used to generate multi-feature maps (MFMs) that annotated ∼94% of the tissue area in DS1 images, with the remaining area comprising alveolar airspace or tissue edges (**Fig. 3b**).

**Figure 3.**
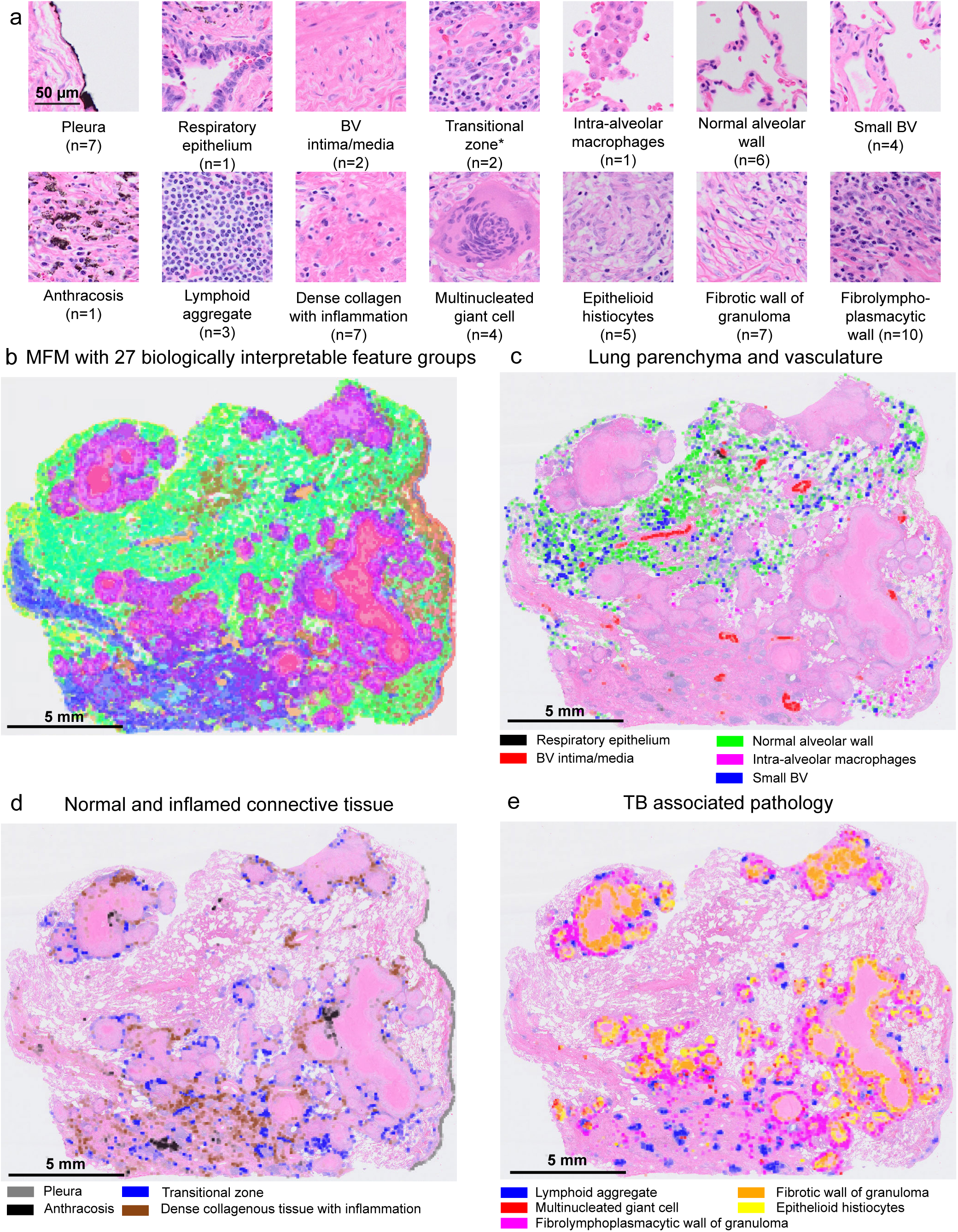
Systematic interpretation of all SAE features in tissue context. **(a)** Exemplar H&E image patches for selected SAE feature groups. The number of SAE features in the group are indicated. Asterisk: transitional zone between disease-involved and normal lung. **(b)** Multi-feature map (MFM), representative H&E WSI with projected score of 27 biologically interpretable groups of SAE1 features (groups 1-27, see **Supplementary** Fig. 4). **(c) - (e)** MFMs on the same WSI with selected feature groups to highlight histomorphologies associated with **(c)** normal lung parenchyma and vasculature, **(d)** normal and inflamed connective tissue, and **(e)** TB-related pathology.

Features associated with normal lung architecture included the normal alveolar wall, respiratory epithelium, various blood vessel (BV) components (BV intima/media, small BV), and intra-alveolar macrophages (**Fig. 3c**). Stromal and connective tissue features captured structural alterations and transitional pathology, including anthracosis (pigment deposits likely due to smoking or air pollution), lung fibrosis manifesting as dense collagenous tissue with inflammation, and the transitional zone between disease-involved and normal lung with reactive pneumocytes (**Fig. 3d**). TB-relevant pathology and immune features included lymphoid aggregates, multinucleated giant cells, epithelioid histiocytes, and distinct layers of the granuloma, including the fibrotic and fibrolymphoplasmacytic (fibrosis, lymphocytes, and plasma cells) walls, as well as necrotic cores (**Fig. 3e**). Because individual patches contained mixtures of histomorphologies (as discussed above), many patches were described by multiple features; accordingly, patch-resolution MFMs commonly associated regions of tissue with overlapping annotations (**Supplementary Fig. 5**). We describe below how to deconvolve and increase the resolution of these annotations.

To verify the reproducibility of MFMs, we assessed SAE feature-score distributions (50th and 90th percentiles) across 40 sequential sections of DS1. Feature scores were similar across sections, showing that the model captures spatially continuous tissue architecture without substantial slide-to-slide technical variation (**Supplementary Fig. 6**). Moreover, when variation was observed, SMEs confirmed that it reflected bona fide differences in tissue architecture across the specimen.

### Alternative sparse autoencoder architectures yield similar histomorphologic feature representations

New approaches to training SAEs are continuously being developed; one recent approach uses a matching pursuit algorithm, which finds optimal projections of multidimensional data, i.e., FM embeddings, onto the elements of a redundant dictionary (i.e., histological features)^56^. This architecture has been reported to exhibit improved reconstruction fidelity compared to standard SAEs, particularly with complex input features. When implemented on UNI embeddings of DS1, the matching-pursuit SAE (SAE2) defined 211 active features. Interestingly, SAE2 suppresses low complexity histomorphologies: only 27% of 211 active features from SAE2 corresponded to necrotic tissue (type 2) or tissue boundaries (type 3) (**Supplemental Fig. 7a**), compared to 70% of 361 for SAE1. Combining all features from SAE1 and SAE2 and clustering by feature overlap generated clusters that usually contained features from both models (**Supplementary Fig. 7b, c**), demonstrating a similar ability to extract the spectrum of histomorphology features. While this may represent a useful advance in future applications of FM-SAE modeling, we used SAE1 for the remainder of this study.

### Multiplexed immunofluorescence independently validates FM-SAE histomorphologic features

To complement SME annotation, we used highly multiplexed immunofluorescence (CyCIF) imaging of DS1 sections adjacent to H&E images to provide independent ground truth of cellular composition (**Supplementary Fig. 2d**). The 39-plex CyCIF antibody panel was designed to characterize parenchymal and immune cell types^45^. Overall, we observed extensive visual correspondence between molecular markers with specific FM-SAE features identified in normal and reactive lungs. For example, large blood vessels (F6261) comprised endothelial cells positive for the transmembrane glycoprotein CD31, surrounded by smooth muscle cells positive for α-smooth muscle actin (α-SMA; **Fig. 4a**). Airway epithelium (F7815) stained positive for pan-cytokeratin (panCK) in columnar epithelial cells and α-SMA in the underlying smooth muscle (**Fig. 4b**). The lymphoid aggregate feature F59985 comprised a mixture of cells positive for the T cell marker CD3E, B cell marker CD20, and follicular dendritic cell marker CD21, consistent with tertiary lymphoid structures (TLS), which form in peripheral tissues in response to chronic inflammation (**Fig. 4c**)^57^. The clustered intra-alveolar macrophage feature (F47380) was characterized by CD206-positive macrophages in airspaces bounded by panCK-positive epithelial cells of the alveolar wall (**Fig. 4d**).

**Figure 4.**
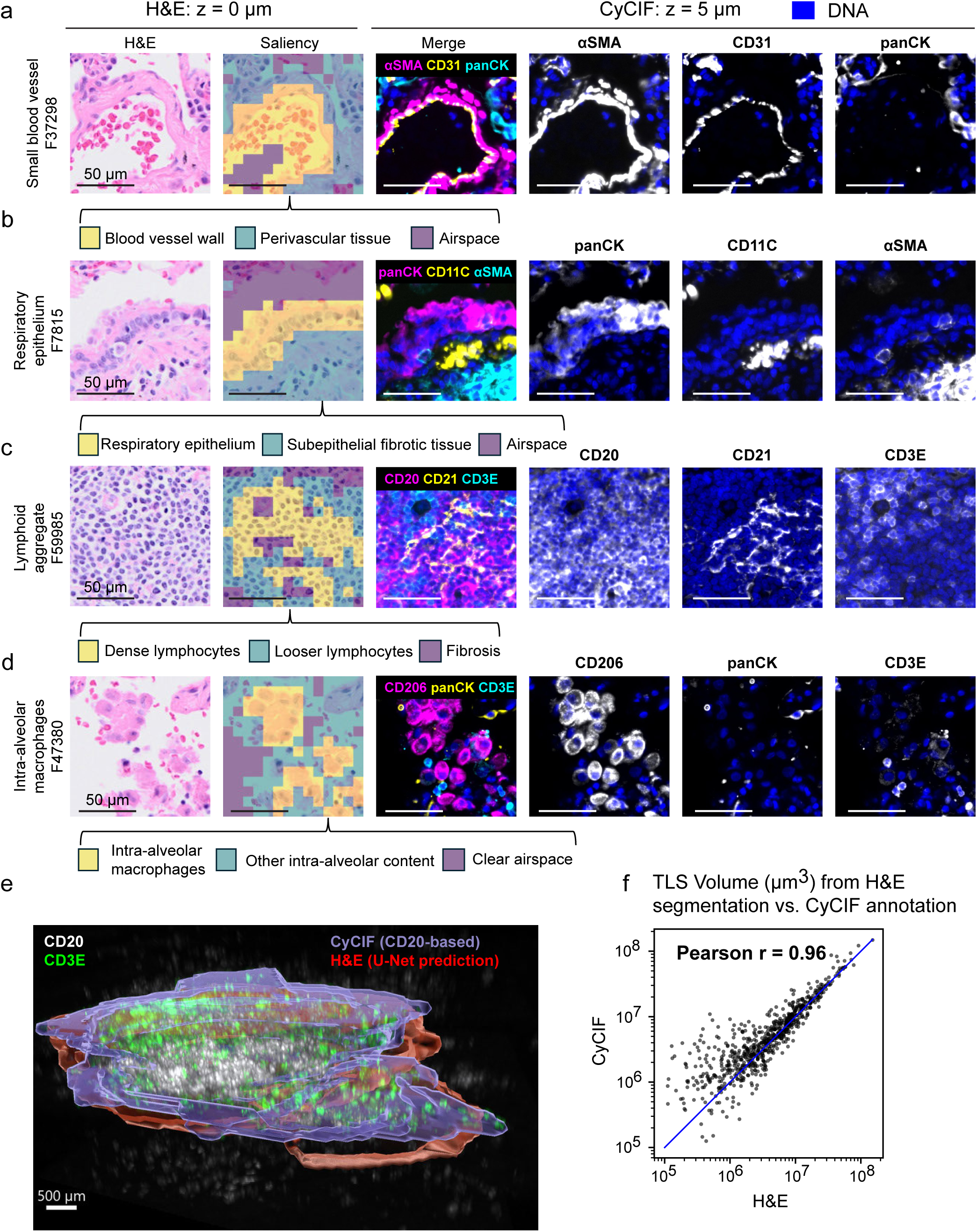
Validating SAE features with multiplexed immunofluorescence (mIF) Exemplar patches of four distinct SAE features. **(a) – (d)** In each row, left: H&E and feature-specific saliency (K=3) map, with interpreted token label (below); right: CyCIF multi-channel merge of adjacent 5 µm tissue section with feature-specific markers (top), followed by DNA (blue) merged with single-marker IF (grayscale) channel for each other marker. **(e)** Three-dimensional rendering of a reconstructed TLS object. CyCIF staining of CD20 (gray) and CD3E (green) in tissues. Boundary of a TLS defined by CD20 threshold (blue) or segmentation mask from H&E-trained U-Net (red), as indicated. **(f)** Scatterplot, LA volumes measured by the two modalities, correlation (Pearson: 0.96).

As a more systematic evaluation of the correspondence between FM-SAE annotations and CyCIF data, we leveraged a recent 3D reconstruction of 420 intact TLS in CyCIF sections adjacent to DS1^45^. In this dataset, TLSs were identified in 3D based on the prevalence of CD20-positive B cells and CD3-positive T cells (**Fig. 4e**, blue envelope). Lymphoid aggregates were identified in H&E images using saliency maps to generate a complementary set of 3D envelopes (red), which were then non-rigidly registered to adjacent CyCIF images using DeeperHistReg^58^. The correlation between TLS volumes derived from CyCIF and H&E data was r = 0.96, indicating a high level of agreement (**Fig. 4f**). Inspection of areas of disagreement revealed that these occurred primarily in regions of relatively low lymphocyte density, where SMEs were themselves uncertain about the precise boundary of the structure (**Supplementary Note**). Together, these results indicate that FM-SAE-derived lymphoid aggregate annotations closely correspond to independent molecular assessments of the same features.

### FM-SAE features capture structural and microenvironmental heterogeneity of multinucleated giant cells

To explore the interplay between SAE-FM-derived features and SME interpretation of these features, we examined multinucleated giant cells (MNGCs)^59^, which are thought to have an important role in phagocytosis of *Mycobacterium tuberculosis* (Mtb) bacilli. MNGCs are prototypical of mesoscale structures that are difficult to identify and study using existing molecular spatial profiling methods since segmentation presumes one nucleus per cell. The large distance between MNGC nuclei and the plasma membrane also complicates phenotyping with cell surface markers. Finally, variability in MNGC morphology makes systematic annotation by humans laborious.

Following *Mtb* infection, activated macrophages differentiate into cytokine-secreting epithelioid histiocytes, which fuse to form MNGCs^59^. This process recruits other immune cells, eventually contributing to granuloma formation (although the precise sequence of events remains uncertain)^60^. The MNGCs most commonly associated with TB are of the Langhans type and appear as an irregular ovaloid cell with a diameter of ∼15 – 120 µm (in 2D sections) containing dozens of nuclei arranged in a horseshoe pattern along the cell periphery. Four FM-SAE features (F5196, F56436, F46327, and F61987) detected giant cells but with substantial morphologic variability across patches and inter-observer disagreement in interpretation. To better understand the origin of this variability, we located MNGC plasma membranes in top-scoring FM-SAE patches and examined patterns of staining for immune and other marker proteins in adjacent CyCIF images. Membrane localization was accomplished at pixel-level resolution in H&E images using saliency maps from ∼1,000 patches to train a convolutional neural network (CNN) based on a ResNet-34 U-Net architecture^61^. **Supplementary Fig. 8a** shows an exemplary MNGC patch and saliency map (F5196; yellow tokens) for a classic Langhans phenotype cell, and a second MNGC with scattered nuclei, surrounded by epithelioid histiocytes (blue tokens). An overlaid U-Net segmentation mask delineates the periphery of the MNGC and corresponds, in adjacent CyCIF images, to the border of mesh-like CD11c staining; the mesh corresponds to the membranes of proximal epithelioid histiocytes (a subset of which were also positive for the macrophage marker CD206). The mask encompasses multiple nuclei circumscribed by the nuclear lamin protein LMNB1, confirming the identification of these cells as MNGCs and validating associated FM-SAE features.

However, comparison of different MNGCs revealed broad variation in H&E and CyCIF morphologies (**Supplementary Fig. 8b**), illustrating the difficulty in consistently characterizing these cells. To better understand how this arises, a single MNGC was imaged across six sequential sections (spanning 75 µm). H&E and CyCIF morphologies were observed to vary substantially with the section (**Fig. 5a**), showing MNGCs to be highly irregular three-dimensional structures and suggesting that a major source of morphological variability arises from differences in where these large cells are transected by 5 µm thick tissue sections. With respect to feature interpretation, we found that higher-scoring patches were more likely to contain large MNGC cross-sections, whereas lower-scoring patches were often centered on adjacent epithelioid histiocytes (**Supplementary Fig. 8c-e).** Together, these observations suggest that variability in FM-SAE feature assignments reflects underlying biological and geometric heterogeneity rather than model instability. Moreover, the combined use of FM-SAE modeling followed by iterative SME annotation provided a computationally accessible representation of complex 3D MNGC structure that extended what was achieved by model or human annotation alone.

**Figure 5.**
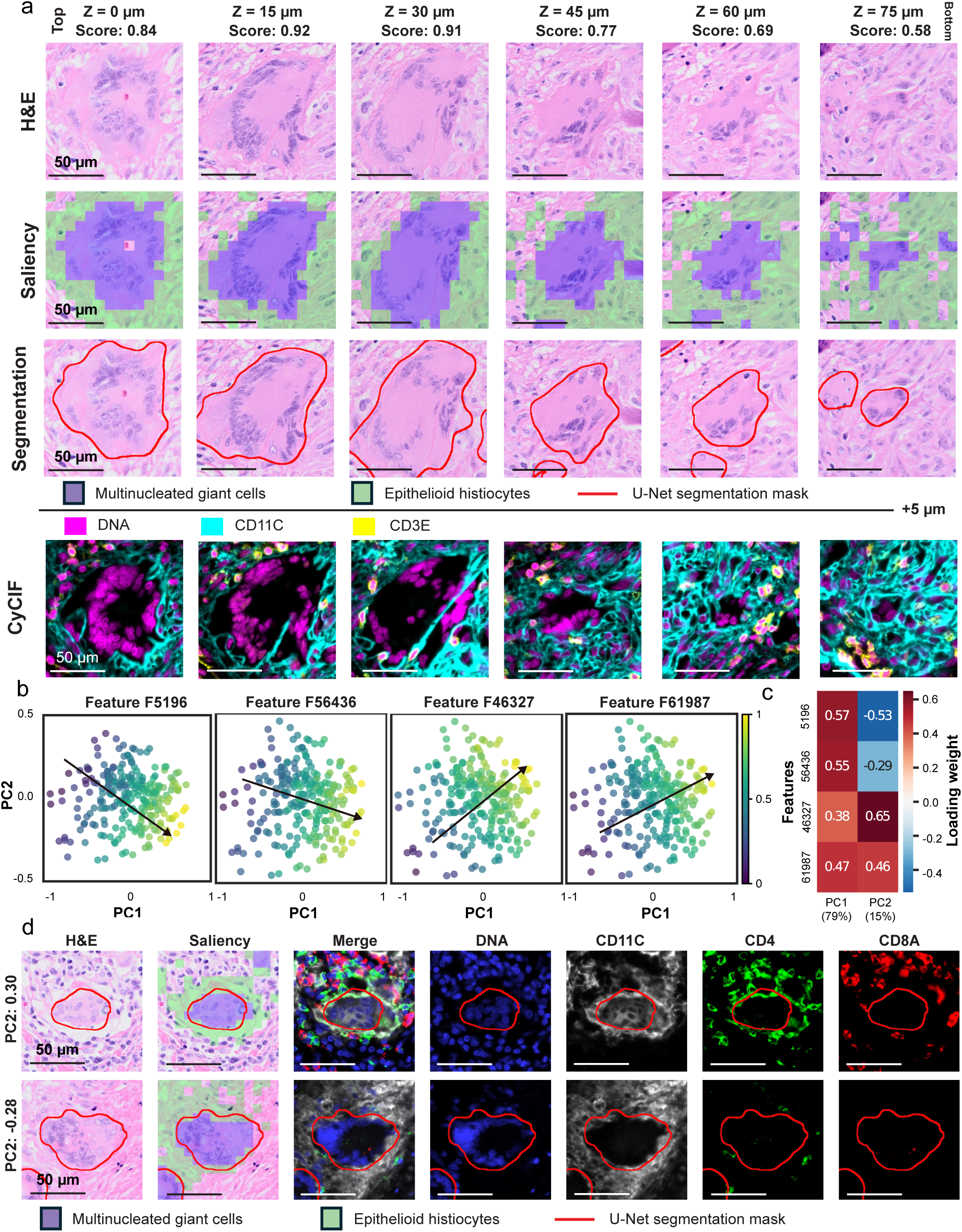
SAE feature recognizes heterogeneous MNGC morphology across serial sections and in different tissue environments. **(a)** Multiple H&E cross-sections of one MNGC (15 µm steps, 75 µm total). Top: Tissue depth and normalized SAE feature score for F5196 (MNGC). Row 1: H&E image; row 2: saliency maps using consolidated token label dictionary and only tokens labeled MNGC (purple) or epithelial histiocytes (green) are shown; row 3: H&E image with boundary (red) of U-Net segmentation of all MNGCs; row 4: CyCIF image merge (z-offset +5 µm): DNA (magenta), CD11C (cyan: MNGC and epithelioid histiocytes), CD3E (yellow: T-cells). **(b)** PCA projection of UNI embedding for curated MNGC (n=245) image patches. Colors indicate patch-level score for each feature (indicated) from low (blue) to high (yellow) with general trend (arrow). **(c)** Heatmap, SAE feature loadings for PC1 and PC2. Explained variance for each PC indicated below. **(d)** Representative patches with highest and lowest PC2 values, indicated left. Each row (left to right): H&E, saliency map as (a), CyCIF multi-channel merge followed by individual channels for the patch centered on MNGC in adjacent tissue section (z-offset +5 µm).

FM-SAE modeling is designed to generate a redundant feature dictionary. To determine whether the four features associated with MNGCs were truly redundant, we performed principal component analysis (PCA) on feature scores for patches centered on 245 MNGCs segmented by U-Net from one section. In the resulting PCA space, all four FM-SAE features projected along PC1 with similar loadings (**Fig. 5b, c**). PC1 correlated in adjacent-section CyCIF data with cell area and nuclear count, the fundamental characteristics of MNGCs. In contrast, the loadings of F5196 and F56436 along PC2 were opposite in sign from those of F46327 and F61987 and correlated with the peripheral intensity of immune cell markers such as CD11c (dendritic cells), CD4 (helper T cells), and CD20 (B cells) (**Supplementary Fig. 8f, g**). Inspection of representative CyCIF images confirmed that PC2 captured differences in the composition of neighboring immune cells (**Fig. 5d)**. We conclude that the four FM-SAE models correspond to two MNGC microenvironments. Thus, while there is some redundancy in the FM-SAE feature library, distinct features carrying the same histological annotation can capture biologically meaningful differences among patches.

### FM-SAE features generalize across specimens and pulmonary diseases

CPath patch-level classifiers constructed using supervised learning often perform poorly outside the training set. This has been attributed to variation in staining intensity, tissue preparation, and scanning hardware, which can lead to overfitting on processing artifacts rather than generalizable biological features^62,63^. To evaluate how FM-SAE features perform in and out of domain, we applied SAE1 features detecting LA and MNGC in DS1 to images from DS2 to DS5. A supervised baseline was constructed using logistic regression on UNI FM patch embeddings; such “linear probing” is a standard approach for benchmarking FMs in CPath^64^. To train linear probes for LA and MNGC, SMEs sparsely annotated these histomorphologies in 39 of 40 H&E images of DS1; one section was withheld and exhaustively annotated as ground truth. SMEs also selected representative sections from out-of-domain datasets DS2 and DS3 and exhaustively annotated all LAs and MNGCs. Using a small balanced training set (e.g., LA vs. not LA) yielded better performance than using all patches from annotated sections. To robustly estimate linear probe performance, we trained an ensemble of probes on random balanced subsets and report the mean and variance of the evaluation metrics.

Visually, FM-SAE features agreed well with human annotations in DS2-DS5 (**Fig. 6a**, **Supplementary Fig. 9**). Both the FM-SAE feature and linear probes achieved >0.97 area under the ROC curve (AUROC) for LA patch classification (**Supplementary Fig. 10a**). However, this metric is inflated by class imbalance since LAs occupy a small fraction WSIs. We therefore switched to precision–recall curves, which better account for the minority class. In-domain, SAE feature F59985 achieved an average precision (AP) of 0.84, as compared to an average of 0.76 for the linear probes on DS1. On DS2 and DS3, the SAE feature consistently outperformed linear probes. Similar trends were observed for MNGCs (**Fig. 6b**) showing that FM-SAE features generalize more robustly than linear probes on out-of-domain datasets of pulmonary tuberculosis. Looking beyond specific annotations to the broader spectrum of histomorphologies, SAE1 features generated MFMs covering >90% of representative sections in DS3 (**Supplementary Fig. 10b**).

**Figure 6.**
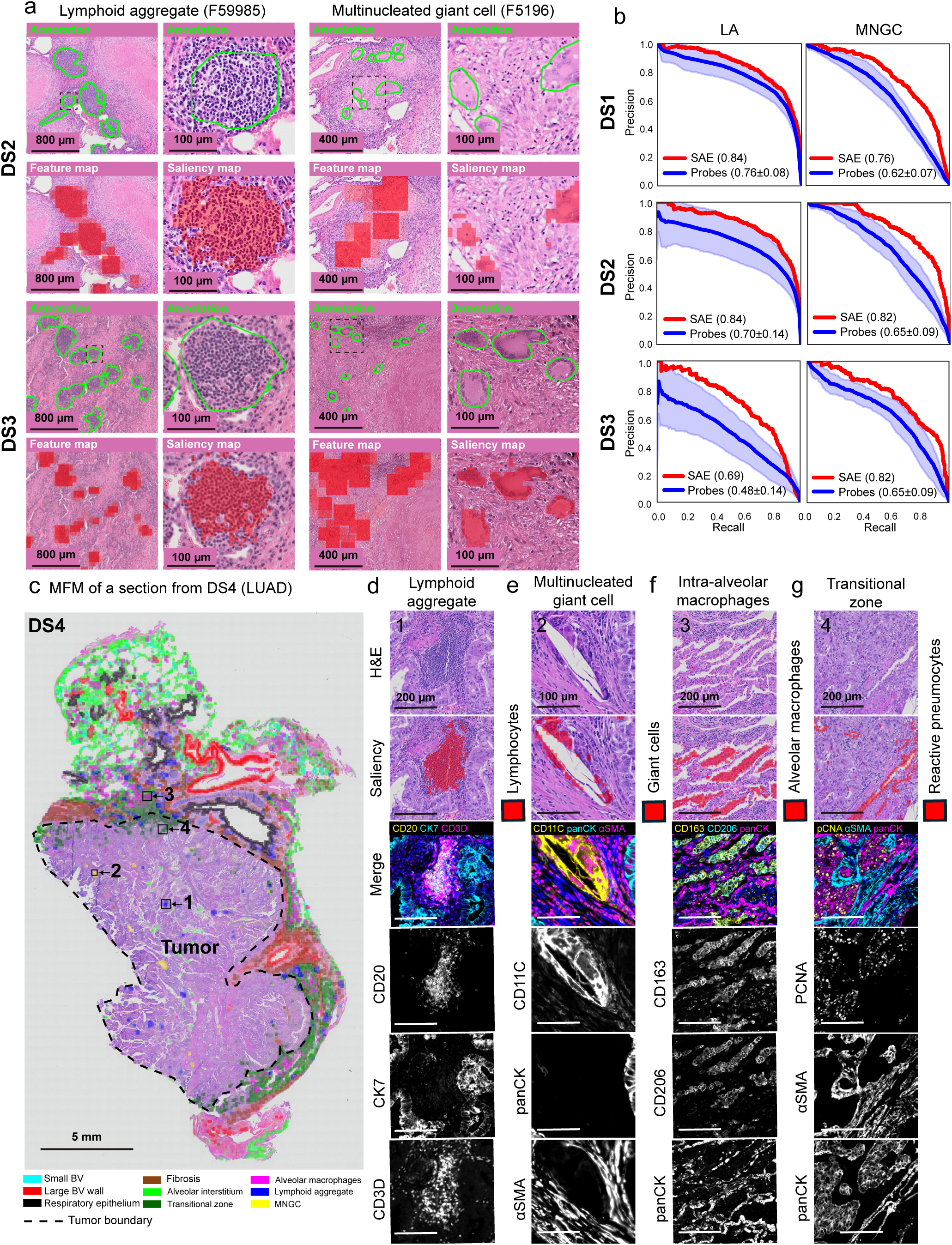
Generalizability of SAE features across datasets. **(a)** Generalizability of F59985 (LA) and F5196 (MNGC) to datasets DS2 (same patient as DS1) and DS3 (different patients). Each quadrant (dataset and feature) includes a representative mid-power (left) and a high-power (right) FOV, focused on area indicated by dashed box (left). Top row, SME-curated feature boundary (green) is indicated. Bottom left: H&E with positive image patches (red). Bottom right: H&E and positive image tokens (red). **(b)** Precision-recall (PR) curves for SAE (red) features versus ensemble of linear probes for LA and MNGC patch-level classification across DS1-3. Ensemble of probes: average (solid blue), range of performance (light blue). For each dataset and feature, average precision (AP) for SAE and ensemble of linear probes (average ± std. dev.), are reported. **(c)** MFM of representative lung adenocarcinoma H&E WSI from DS4. Tumor boundary (dashed line) and color map to features is defined (bottom). FOVs (solid boxes, arrows) 1-4 map to panels (d) – (g), respectively. **(d) – (g)** H&E image (top) from FOV in (c), for indicated histomorphologies (above). Second, H&E with map (red) for indicated token-level saliency (right). Next, CyCIF multi-channel merge, followed by single-channel (gray) images with marker names (left).

To assess generalizability to other diseases, we applied the TB FM-SAE features to non-small cell lung cancer (NSCLC; DS4) and sarcoidosis (DS5). Excluding tumor regions, which were not represented in DS1, MFMs labeled >80% of a typical NSCLC image (**Fig. 6c**). SME inspection, token saliency, and CyCIF imaging all confirmed the presence of common histomorphologies such as LAs in TB and NSLC lungs (**Fig. 6d**). MNGCs were also present in NSCLC, but many were foreign body giant cells (FBGCs) surrounding cholesterol crystals, which are associated with chronic inflammation (**Fig. 6e**). Giant cells were also detected in H&E images of sarcoidosis; these are known to be of the Langhans-type^65^ **(Supplementary Fig. 11a)**. Clusters of intra-alveolar macrophages were also found in association with both tumor (**Fig. 6f)** and normal-appearing epithelium (**Supplementary Fig. 11b**). A “transitional zone” between tumor and normal tissue was also found to be similar in morphology and composition to transitional zones around TB granulomas, pointing to biological similarities that warrant deeper investigation (**Fig. 6g**). We conclude that the ability of TB-derived FM-SAE features to identify related histomorphologies in distinct diseases such as NSCLC and sarcoidosis supports their potential for broad out-of-domain applicability.

## DISCUSSION

This paper makes three primary contributions to the rapidly growing fields of computational pathology and spatial biology. First, it shows that a primary barrier to human interpretation of FM model embeddings is the presence of multiple histopathologies in each of the image patches used by the model. These patches have a spatial scale of 100-400 cells, and intra-patch heterogeneity makes it difficult to interpret patch labels and clusters in biological terms. Second, this challenge can be largely overcome by using a compositional approach (in this work, using SAEs) that learns to assign each patch to multiple features, each corresponding to the different histological component. Interpretability is further enhanced by the availability of continuous feature scores and token-level saliency maps with approximately cell-level granularity within the image patch. Whole-slide views generated in parallel show how each SAE-FM feature maps to macroscopic tissue architecture, recreating how histopathologists examine physical slides at both high and low magnification. Third, once relationships between FM-SAE features and histopathology concepts have been established, it is possible to automatically and systematically annotate in-domain and out-of-domain H&E images and use these images to interpret highly multiplexed tissue images from adjacent tissue sections (and vice versa). We see this as a particularly valuable application: molecular spatial profiles are ideal for single-cell analysis, but they struggle to identify mesoscale tissue features, particularly when those features include acellular components. The patch-level representation of images in FMs is highly advantageous at this larger spatial scale, and their training on H&E images provides a direct connection to fundamental knowledge of histopathology. In contrast, manually annotating large sets of H&E images is time consuming and rate-limiting; it is also uncommon to have access to multiple SMEs for this task.

FMs are trained in a self-supervised manner, and this was also true of SAEs in this manuscript, but interpretation of model outputs was performed by human experts. We found that SMEs agreed almost immediately on the most obvious multicellular structures such as airways, blood vessels, fibrosis, lymphoid aggregates, and granulomas of different types as well as localized features such as anthracotic pigment missed or diluted by the clustering approach. The use of saliency maps at the level of tokens, which in UNI are 196-fold smaller than patches and have the approximate dimensions of a single cell, helped to resolve the semantics of compositional features recovered from embeddings. During this work, we developed provisional annotations not only for patches but also for their component tokens. The availability of next-section high-plex CyCIF data served as an independent source of information on these tokens.

In the case of structures with highly variable appearance, such as MGNCs, we found that FM-SAE models detected structures missed during human “ground truth” annotation, leading to the impression that the FM-SAE model lacked precision (e.g., due to hallucination), but subsequent re-inspection of images by SMEs suggested that the model was correct. The simple interpretation of this finding is that humans tire when performing highly repetitive tasks (e.g., counting many instances of the same structure) and do not create an accurate ground truth (this also shows up as inter-observer variability)^66^. A second interpretation is that observers actually learn by interacting with the FM-SAE model, resulting in better understanding of the diversity of a histological feature. Whether this was true of SMEs remains uncertain, but it was certainly the case for the biologists working on the project. In either case it results in ground truth annotations that are incomplete, necessitating recursive human-in-the-loop evaluation that complicates rigorous model evaluation.

We found that FM-SAE features developed for TB-infected lung collected could be applied to lung diseases such as non-small cell lung cancer. This contrasts with poor out-of-dataset (OOD) performance observed for many CPath models due to domain shift and ascribed to variables such as tissue processing, staining, and scanning^67,68^. We found that the average precision of our FM-SAE model was comparable to supervised linear probes in-distribution while substantially outperforming them under domain shift. This is promising with respect to using FM-SAE annotation at scale, but it will still be necessary to create consistent patch- and token-level feature dictionaries. Ideally, these dictionaries will align with molecular definitions from the Human Cell Atlas^69^ and Human Reference Atlas (HRA; part of HubMAP)^69^ and with histological and anatomical definitions from the Terminologia Histologica^70^, International Federation of Associations of Anatomists^71^ or similar international projects.

### Using SAEs to analyze FM model embeddings

Computational approaches other than SAEs, such as soft clustering, topic modeling, and dictionary learning, may also be able to address the limitations of discrete clustering of FM embeddings. However, SAEs offer multiple advantages. First, by utilizing mini-batch gradient descent during training, SAEs scale efficiently to millions of patches, in contrast to traditional dictionary learning. Second, by projecting FM embeddings into an overcomplete feature space, SAEs can detect rare or subtle morphological variation. Third, the use of SAEs in histology connects to the emerging use of autoencoders to study data representation by large language models in domains such as natural language, protein and genetic sequences, and medical images^55,44,72–74^. The SAE architecture is not new, but continues to evolve; in the current work we explore training via the matching pursuit algorithm^56^; this yielded similar features to a standard approach but may be advantageous in suppressing low complexity features. This argues for further investigation of other SAE architectures.

### Applications in spatial biology and AI-assisted diagnostic histopathology

Is the effort involved in aligning the output of FM models with human understanding worthwhile? In the area of CPath, it is generally accepted that pathologists and diagnostic models will need to work together^75^. Pathology journals also evidence a high level of interest in the transformative potential of spatial biology and deep molecular analysis of tissues. The reciprocal is not as evident: there is an enduring assumption that computational methods can learn biology ab initio from multiomic data. We do not think this likely since human understanding derives from a conceptual understanding of tissues that is based on more than inspection of images. Thus, we envision detailed FM-SAE-based annotation aligned with human understanding as a routine complement to high-plex tissue imaging and spatial transcriptomics. Combining FM-SAE modelling of the same or adjacent section H&E images with combinatorial channel multiplexing^76^ would potentially achieve a highly cost-efficient approach to large datasets. A second consideration is that histological stains are uniquely effective at identifying multicellular structures with distinctive textural features^77^, extracellular matrix^78,79^, and areas of necrosis having few if any cells. In the current work, anthracosis and fibrosis would be missed entirely by profiling methods that rely on gene or protein expression in single cells. MNGCs and other multi-nucleated cells would also be missed because they violate the assumption in standard segmentation algorithms^80–82^ that each cell contains one nucleus. A final consideration is training FMs on patches containing several hundred cells makes them more effective at identifying mesoscale features such as blood vessels than purely single-cell methods.

### Limitations and future research

The patch- and token-level labels used in this work have been reviewed by SMEs but remain a work in progress. SAEs were trained without supervision, and interpretation was added only after the fact, but future models could be improved by incorporating biological priors that down-sample low-complexity (or non-informative) morphologies or utilize hierarchical or structured sparse representations^83,84^. We consciously focused on analysis of H&E models alone, using multiplexed imaging as a second source of ground truth, because H&E images are numerous in the literature and the primary source of histological prior knowledge. However, one extension of FM-SAE modeling will be to integrate learning on image-associated text and multiplexed data. Moreover, while unsupervised discovery is powerful on its own, SAE feature scores could also be calibrated with a small set of expert curated labels to help identify features of particular importance (for example, potentially distinguishing arterioles from venules). Such an effort would likely go hand-in-hand with the development of standardized patch and token-level feature dictionaries.

## ACKNOWLEDGMENTS

This work was primarily funded by a grant from the Bill and Melinda Gates Foundation INV-027106 with additional support for study of cancer tissues from an ASPIRE award from the Mark Foundation for Cancer Research, an NIH grant (R24AI186591; A.S.), and the Wellcome Trust Core award: 227167/A/23/Z of the Africa Health Research Institute. We thank the Neurobiology Imaging Facility (NIF) at Harvard Medical School for imaging support and members of the LSP spatial profiling team, particularly B.K. Kobs, J. Muhlich, M. Lopez Leon and Y. Chen for expert assistance throughout the project.

## OUTSIDE ACTIVITIES

P.K.S. is a cofounder and member of the Board of Directors of Glencoe Software and a member of the Scientific Advisory Board for RareCyte and Montai Health; he holds equity in Glencoe and RareCyte.

P.K.S. is a consultant for Merck and Danaher.

## AUTHOR CONTRIBUTION STATEMENTS

Z.Z. and Z.M. conceived and designed the study. A.S., S.S. T.N. and P.K.S. co-supervised the work and acquired funding. Z.Z. developed and implemented the data analysis workflow and produced the figures with input from Z.M., P.K.S. and other authors. Z.Z. and Z.M. interpreted the results guided by the expertise of SMEs A.G., T.N., and S.C. I.H.S and A.S. acquired specimens, and E.O and Z.M. acquired H&E and CyCIF images, and Z.M., E.O., S.R.T. K.L. performed image analysis. Z.Z., Z.M. T.N. and P.K.S. wrote the manuscript with input from all authors.

## Code availability

Original code associated with this Article is available via GitHub at https://github.com/labsyspharm/Zhao-FMSAE-2026.

## ONLINE METHODS

### Sample acquisition and dataset preprocessing

Formaldehyde-fixed paraffin-embedded (FFPE) tissue samples were obtained from biopsies and autopsies archived at Brigham and Women’s Hospital (BWH) in Boston, Massachusetts (United States of America), and Africa Health Research Institute (AHRI), University of KwaZulu-Natal, Durban (South Africa). Tissues were screened by acid-fast staining for *Mycobacterium tuberculosis.* Specimens were collected under BWH IRB 2018P001627 and the University of KwaZulu-Natal Biomedical Research and Ethics Committee (BREC) protocol BE019/13. Sectioning and H&E staining of FFPE tissues were performed at BWH or AHRI (**Supplementary Table 1**).

For DS1, 120 5-µm thick adjacent tissue sections were prepared from the ∼2 x 1.5 cm tissue block from a treatment-naïve patient (Patient 1_B4) previously assessed to have both normal lung architecture and large necrotizing and non-necrotizing cellular granulomas. Sections were cut onto Superfrost Plus (Thermo-Fisher) histopathology slides for subsequent processing. 40 H&E and 40 CyCIF images were obtained, skipping one interleaving section to capture representative tissue morphology across 600 µm tissue depth. Histopathological review and annotation of a subset of the H&E-stained serial sections was performed by a board-certified pathologist using OMERO Pathviewer (Glencoe Software), a web-based image browser.

H&E-stained FFPE sections for DS1 and DS2 were scanned on an Olympus VS200 automated slide scanner equipped with a 20X objective (NA 0.8; 0.2738 µm/pixel) at the Neurobiology Imaging Facility at Harvard Medical School. Imaging for DS3 was performed at AHRI using a NanoZoomer S60 digital slide scanner (Hamamatsu Photonics) in brightfield mode using a 20x objective (NA 0.75) and 2x magnification changer to yield a final spatial resolution of 0.23 µm/pixel.

Multiplexed CyCIF imaging was performed on FFPE tissue sections, as previously described^85^ and is described in full in Ogbonna et al^45^.

### Tissue segmentation and patching

For each whole slide H&E image (WSI), tissue area was first segmented and cropped to generate a library of overlapping tissue patches. To generate a tissue mask, WSI was downsampled and transformed from RGB to HSV color space for entropy guided image segmentation. The local entropy of pixel intensity distribution was calculated for the saturation channel, followed by a median filter to smooth the output and, finally, Otsu thresholding to separate high (tissue) and low (background) entropy regions, generating a binary tissue mask^86^. For lung samples, small holes in this mask were filled to fully sample regions of respiratory ducts and alveoli. Overlapping image patches of size 448 pixels (∼120 µm) in x and y dimensions were cropped from regions within the tissue mask using a regular grid with spacing of 256 pixels. This sampling gives a 43% overlap in each dimension between adjacent patches, ensuring complex histomorphologies are adequately represented. Only patches with centroids inside the tissue segmentation mask were kept for the next step of processing.

### Unsupervised histomorphology feature discovery with SAE

All image patches from 40 H&E sections in DS1 were encoded using the UNI foundation model to produce, for each patch, a 1,024-dimensional embedding to capture the textural variation^26^. A sparse autoencoder (SAE) with a single linear layer followed by ReLU activation and a linear decoder was trained using all patch embeddings^55,87^. To provide an overcomplete representation of the UNI embeddings, we utilized a latent dimension of 65,536 to ensure the model captures rare or subtle morphological variations that might otherwise be lost in the lower-dimensional bottlenecks characteristic of traditional clustering. The loss function consists of a reconstruction term, which encourages the decoder output to be similar to the input embedding to the encoder, and an L0 regularization that encourages the reconstruction to use few features. All patch embeddings were first normalized by rescaling each embedding dimension to have a unit root mean square value, and the same normalization factor was a model parameter kept the same when processing other datasets. The model was optimized by gradient descent with an Adam optimizer for 200k steps, with a learning rate of 5 × 10^−5^ and a batch size of 2,048. The weight on L0 regularization was dynamically adjusted every 10 steps to approach a target sparsity of 0.001 (0.1% SAE features out of 65,536 active), which was empirically found to be stable across several other datasets with 200k up to 3M patches.

After training, for each SAE feature, both the maximum feature score and the percentage of all patches with nonzero score were computed. Features with low scores or infrequent activation were discarded, resulting in a total of 361 features discovered by the model. We noted that for different datasets, the target sparsity may require tuning to reach similar numbers of active features. At L0 sparsity loss of 20.69 (relative to 65,536), for SAE1 we obtained an L2 reconstruction loss of 0.067 and a fraction of variance unexplained (FVU) of 0.069. After filtering, the remaining SAE1 features achieve an L0 sparsity of 15.60 and the same L2 reconstruction loss and FVU, suggesting the most important features are retained without sacrificing reconstruction fidelity.

SAE2 uses the recently developed MP-SAE architecture, where, by unrolling the matching pursuit algorithm, the encoder layers sequentially select an approximately orthogonal feature to reconstruct the residual from previous layers^88^. SAE2 was initialized with a latent dimension of 8,192 and iteratively reconstructs the input embedding up to a threshold of 0.01 in Euclidean distance. The model is trained with the Adam optimizer, with a learning rate of 0.03, up to 50,000 steps with a batch size of 8192.

The FM embeddings were extracted, and the SAE1 and SAE2 were trained using PyTorch (v2.6.0) on the HMS O2 high-performance computing platform, using 32GB RAM, 4 CPU cores (Intel(R) Xeon(R) Platinum 8468) and 1 GPU core (RTX 8000 with 48GB VRAM, CUDA version 12.8).

### Generating token-level saliency maps and feature reports for SME review

PDF feature reports for SME review were generated using the *spatialdata_plot* package. For baseline comparisons, histomorphologic features are defined as clusters from discrete clustering methods: K-means with K=25 and Leiden algorithm using default settings for *neighbors* and *leiden* functions implemented by the *scanpy* package. The feature reports included 20 randomly sampled image patches assigned to the respective cluster, which were then projected as colored regions onto 10 representative whole-slide images (WSIs; sampled as every fourth WSI in DS1).

For the 361 active SAE features, each report included 10 image patches randomly selected from the top-scoring decile (10%). To facilitate SME interpretation of intra-patch heterogeneity, we generated feature-specific token-level saliency maps for these top-scoring patches. To construct these maps, 1,000 patches were chosen evenly from the top decile as positive examples, alongside 1,000 randomly selected zero-scoring patches as negative examples. Positive examples were embedded using the foundation model without global pooling to extract 14×14 token-level embeddings. A K-means model (K=3) was then fit to these token embeddings.

For any given image patch in the report, its feature-specific saliency map was produced by extracting the token embeddings, applying the trained K-means classifier, and up-sampling the cluster labels to match the patch size. These token clusters were visualized by shading the tokens in the 10 representative H&E image patches, with shading intensity determined by the relative enrichment of token labels in top-scoring versus zero-scoring patches (highly enriched clusters were assigned a lighter color for visual emphasis). Finally, continuous SAE scores were projected onto the 10 representative WSIs to produce macroscopic WSI feature maps.

### Aggregating SME notes for a consolidated dictionary of token labels

Feature scores were first normalized for each feature across DS1 to 0-1 range. Pairwise feature overlap was computed as set level IoU, defined as the fraction of patches with high scores (>0.5) for both features (intersection) over the patches with high scores for either (union). The overlap matrix was used as a distance matrix and hierarchically clustered with average linkage to reorder the features for plotting. Based on reviewing the feature overlap heatmap and morphology and individual feature reports, SMEs grouped features, resulting in 105 cellular features in 29 groups (including one group for blank space or debris and one group for miscellaneous features, besides 27 biologically interpretable groups). SMEs also interpreted feature-specific saliency maps for at least one feature in each group. As multiple features, sometimes even from different groups, might be given the same token level label, we aggregated the SME notes to group similar token clusters together. This was accomplished by hierarchical clustering of centroids pooled from all K-means (K=3) from different SAE features, using the cosine distance metric. The dendrogram was then plotted with feature group index and SME interpretation on each leaf. We manually merged dendrogram branches with similar SME annotations (distance threshold = 0.3), consolidating them into 18 distinct token-level labels. Using the centroids of these labels, we built a 1-nearest neighbor model to generate comprehensive saliency maps encompassing all 18 labels present in DS1, expanding upon the initial K=3 feature-specific maps.

### Generating multifeatured maps (MFM) to capture tissue organization

Scores of features from each group were aggregated by taking the maximum normalized scores over the group, which achieves better coverage for histomorphologies that have variable presentations deemed equivalent by SMEs. To generate multi-feature maps (MFMs), spatial overlaps between image patches were resolved using a custom spatial max aggregation function optimized for sparse feature maps. Specifically, positive-scoring patches were projected onto the WSI, retaining only the highest patch-level feature score in any overlapping region.

### Pixel-level U-Net segmentation of tissue morphologies

The SAE1 feature F61987 was used to generate a U-Net classifier for MNGCs in DS1 H&E images. We 1,000 top patches and 1,000 zero-scoring patches as positive and negative samples to form the training dataset. We obtained token-level saliency maps as previously described. The MNGC cluster, which was visually reviewed by experts, was selected to binarize the token-level map. This cluster denotes the segmentation target, and the other two are background. The binarized mask was then smoothed with a Gaussian filter to form a per-patch ground truth mask, and the negative patches have empty masks. We employed a U-Net architecture with a ResNet-34^89^ backbone, trained using DICE loss. Optimization was performed using the Adam optimizer with an initial learning rate of 2 × 10^−4^, governed by a cosine annealing scheduler (minimum learning rate of 1 × 10^−5^). The model was trained for 10 epochs with a batch size of 32 on the balanced positive and negative examples. SAE1 features F27188 and F59985 were used to develop a pixel-level classifier for LAs in DS1 similarly, using a doubled patch size to adequately capture the tissue context. The trained U-Net was then applied to large, overlapping FOVs across each WSI. Predicted probabilities were averaged in overlapping regions to avoid boundary artifacts, and a threshold of 0.5 was applied to binarize the prediction into a segmentation mask for MNGCs and LAs.

### H&E to CyCIF image registration

DeeperHistReg^58^ was used to register each H&E to the first DNA channel of the adjacent CyCIF WSI in two steps: an affine transform for crude image alignment followed by a deformation field to match local texture features. The combined transform was stored by DeeperHistReg as a 2D array of displacement vectors that, in the properly shifted coordinate system, describes how each point in the target image (CyCIF) maps back to the source image (H&E). The deformation vector field was numerically inverted to map image patch centroids from H&E onto CyCIF images.

### TLS label transfer from CyCIF to H&E

Ogbonna et al., 2026, reconstructed 3D TLSs from serial CyCIF sections following ANTs-based image registration^45^. Connected-component analysis of binarized 2D TLS ROI masks in MATLAB (R2022a) was then used to generate instance-labeled 3D TLS annotation masks. To facilitate comparison of H&E with the CyCIF-based annotations, raw whole-slide segmentation mask from the previous section was registered to the DNA1 channel of each adjacent-section CyCIF from the registered 3D stack of CyCIF images, resulting in an initial 3D segmentation mask based on H&E. Next, connected components (corresponding to LA structures) of this 3D mask were labeled, and after inspecting and thresholding the histogram distribution of component sizes, small, connected components due to segmentation or registration artifacts were discarded. We then use the labeled regions from the CyCIF mask as seeds for watershed algorithm to transfer and expand the annotations within the H&E mask. Connected components from H&E disjoint from the CyCIF mask were assigned unique labels from the latter for future analyses.

### Characterizing the heterogeneity of MNGCs with SAE features

We applied the trained U-Net to LSP24530 of DS1 to produce a raw, whole-slide segmentation mask and registered it to the adjacent CyCIF image. As with LA, the mask is labeled and small connected components were discarded. Manual QC was done to eliminate labeled instances that contain more than one MNGCs tightly packed together, too small, or had visible local registration error compared to CD11c staining, resulting in 245 labeled MNGCs.

Patches of the same size were placed over MNGCs, and the four feature scores were computed. For cell (nuclei) segmentation, we used Cellpose4^90^ on the H&E patches directly, and while, as expected, the model cannot segment the entire MNGC in the patch, it segments the multitudes of nuclei inside rather accurately. The area and nuclei counts are computed from the area of labeled MNGC region and the count of nuclei segmentation centroids in the region, respectively. For each CyCIF marker, the peripheral staining intensity was computed by averaging signal within the annular region extending ∼10 µm from the boundary of labeled MNGC region on both sides.

### Evaluating the performance of SAE features across datasets

Following the convention in evaluating other histopathology foundation models, we formulate the LA and MNGC detection tasks as patch-level binary classification, i.e., given an image patch, whether the patch contains MNGC or LA. To train linear probes as the supervised baseline, SMEs sparsely annotated representative LAs and MNGCs on WSIs for the remaining 39 out of 40 sections. For evaluation, the first section from DS1, and one section from DS2 are chosen as representative sections; for DS3, we had to choose separate representative sections for LA and MNGC because none abundantly features both. All LAs and MNGCs in these representative sections are thoroughly annotated by SMEs.

The supervised linear probe is implemented as a logistic regression model on top of the patch-level FM embedding. For the classification task, we construct datasets from whole slide images by taking patches and then assigning a binary label as ground truth based on whether the area of the patch covered by any annotation exceeds thresholds set for LAs (0.5) and MNGCs (0.1). The training datasets for the linear probes are constructed by including an equal number of positive and negative patches, whereas the test datasets include all patches from tissue areas across the slides.

To quantify the performance of SAE features and supervised linear probes, we chose both average precision (AP), which is the area under the precision-recall curve, and area under the receiver operating curve (AUROC), with both curves computed by varying the decision threshold over SAE feature scores or predicted probabilities from the linear probes, using the Python package *sklearn*.

## SUPPLEMENTARY FIGURE LEGENDS

**Supplementary Figure 1. Defining histomorphologies by clustering FM patch embeddings**

**(a)** UMAP, sub-sampled (n=50,000) patch embeddings from UNI FM. Color represents patch assignment to K-means clusters (K=25). **(b)** Multi-feature maps for a representative tissue section using K-means with colors as in **(a)**. **(c)** UMAP for the same sub-sampled patch set. Color represents patch assignment to Leiden clusters (n=21). **(d)** Multi-feature maps the same section using Leiden algorithm with colors as in **(c)**. **(e)** Feature report for cluster C17. Top, 20 representative H&E image patches with source WSI indicated, bottom, 10 WSI H&E feature maps with x and y pixel positions indicated. **(f)** Representative image patches for K-means clusters classified based on morphology as class 1: cellular structure, class 2: homogenous, acellular material, class 3: blank space and debris. Right, cluster indices and interpretations shown. **(g)** Selected representative H&E image patches from C17 (lymphoid aggregate, LA) and C7 (fibrosis, F) annotated for regions of F or LA, with boundary indicated (dashed). LA/F indicates FOV with no clear boundary.

**Supplementary Figure 2. Workflow for model training and human interpretation**

**(a)** Initial dataset preprocessing and model training. H&E WSIs are tiled into overlapping patches and embedded using a FM (UNI). A one-layer SAE is trained to reconstruct the FM embeddings using sparse features. Only active features are kept (**Supplementary Fig. 3a**, methods). After training, all FM embeddings are encoded by the trained model into SAE features. Right: each row is the FM embedding of one image patch and each column is an SAE feature. For any patch, most features are not active because of the feature sparsity imposed during model training. **(b)** For each pair of SAE feature, we compute feature overlap as the fraction of patches with high score for either feature that is high for both. The features are ordered by hierarchically clustering the spatial overlap matrix. Combined with feature reports (**Supplementary Fig. 3d**), SMEs could rapidly filter out features detecting blank space, debris, or acellular morphology. **(c)** For each SAE feature, patches are ranked by feature score, and high scoring patches are again embedded by the FM to produce token-level embeddings. Token-level clusters are mapped back to the patches to generate saliency maps to be included in feature reports. Most features are interpreted by this multiscale approach and grouped together, leading to multi-feature maps for whole-slide annotation. **(d)** For a particular feature of interest, a U-Net is trained to produce pixel segmentation that refines the patch level feature score by predicting relevant regions indicated by saliency maps. The trained U-Net is then applied across the whole slide, and the segmentation is transferred to adjacent section subjected to spatial profiling, such as with CyCIF, to validate the molecular morphology of the detected feature.

**Supplementary Figure 3. SAE model training and evaluation**

**(a)** Scatterplot, maximum feature score and fraction of active patches for all primary SAE features (n=65536). Features in the top right quadrant are considered active (orange). **(b)** Spatial overlap matrix for all 361 active features from SAE1 sorted by hierarchical clustering. Color bar (bottom) indicates visual classification of exemplar H&E image patches. **(c)** Montage of exemplar H&E image patches for all 361 active features from SAE1, as ordered in (b). Primary feature index (0-65535), right of each image. Color bar (bottom) indicates classification of features, as (b). **(d)** SAE feature report for F27188. Top, 10 highest scoring H&E image patches and, below, corresponding token-level saliency maps for K-means (K=3) clustered of FM embeddings. Bottom, 10 representative feature map, image patch scores projected by intensity (red). Source WSI for H&E image patches, feature maps, and x and y pixel coordinates indicated on axes of feature maps, as indicated.

**Supplementary Figure 4. SAE feature grouping and consolidated token labels in DS1**

Exemplar H&E image patches (above) representing 29 groups of SAE features, and saliency maps (below) for token-level mapping of 18 distinct token groups. Left: feature group index, in order left-to-right) and named histomorphology. Color coding and SME interpretation of token groups, as indicated (top).

**Supplementary Figure 5. SAE features deconvolve tissue regions with overlapping histomorphologies**

H&E and corresponding MFM of FOV with multiple histomorphologies. Each feature map in MFM is binarized and has boundary colored solid to delineate regions that can overlap.

**Supplementary Figure 6. Consistent feature representation in multiple sections of the same tissue block**

For each SAE feature group, the median (blue) and 90^th^ percentile (orange) of aggregated feature score is plotted across 40 consecutive sections from the tissue block.

**Supplementary Figure 7. Alternative SAE algorithm provides broadly similar features**

**(a)** Montage, H&E image patch exemplars of all SAE2 features. Image border color coded by clustering feature overlap similarly as in **Fig. 1c**. **(b)** Spatial overlap matrix of cellular SAE1 (n=110 out of 361) and SAE2 (n=153 out of 211) features grouped into 25 clusters by hierarchical clustering using complete linkage. Color bar (right or top) indicates feature from SAE1 (green) or SAE2 (red). **(c)** Frequency of cellular SAE1 and SAE2 features in each of the three types of clusters, depending on whether the cluster contain only SAE1 features, only SAE2 features, or a mixture of SAE1 and SAE2 features.

**Supplementary Figure 8. Diversity of MNGCs in H&E images**

**(a)** Exemplar H&E image patch for feature F5196 (MNGC). Left to right: H&E image, feature-specific saliency map (K=3), multi-channel merge, and individual channels are shown on the same row. **(b)** Top row, three representative H&E patches selected of high, intermediate, and low F5196 feature score; middle: saliency map with consolidated dictionary, MNGC (violet) and epithelioid histiocytes (green); bottom: CD11c staining in registered, adjacent section (z-offset +5 µm). U-Net derived MNGC segmentation boundary is indicated (red) in all images. **(c)** Precision versus feature score for F5196. Arrows are color-coded to match (b) to illustrate top, medium, and low scoring patches. **(d)** Representative patches with low score for F5196 without MNGCs. The H&E patch, saliency map, and CD11c staining are shown in columns similarly to (b). **(e)** Scatterplot, correlation (Spearman: 0.74), of feature score and fraction of MNGC tokens. **(f)** Spearman correlation of PC1, PC2 and spatial features of U-Net segmented MNGCs in adjacent CyCIF image: cell area (red), nuclei within MNGC mask (purple) and mean fluorescent intensity for CD4, CD11C and CD20 at mask periphery (light blue). **(g)** Scatterplot, PC2 vs. CyCIF feature from (f) for MNGC-centered image patches. Spearman’s rank correlation coefficient and p-value as indicated.

**Supplementary Figure 9. Accurate feature recall in multiple datasets**

For each combination of dataset from DS1 – 3 and histomorphologies from LA and MNGC, the whole-slide H&E image of the representative section is shown with areas annotated by SMEs in green and SAE feature map in red. SAE feature indices in figure legend.

**Supplementary Figure 10. Extended results demonstrating the generalizability of SAE features in other TB specimens**

**(a)** Receiver operating characteristics (ROC) curves for SAE features (red) versus ensemble of linear probes (blue) for patch-level LA and MNGC classification in DS1 – 3. Average (solid line) and range (light zone). In parentheses, area under ROC curve (AUROC) for SAE and ensemble (average ± std. dev.). **(b)** MFMs, two representative H&E WSIs from DS4 projected score of 27 biologically interpretable groups of SAE1 features (same as **Fig. 3a**).

**Supplementary Figure 11. Extended results demonstrating the generalizability of SAE features in other lung disease specimens**

**(a)** FOV, alveolar macrophage (F47380) in DS4 (LUAD) similar to DS1. Left to right, H&E image patch, token level saliency map (red, consolidated dictionary), multi-channel merge, and single-channel grayscale images. **(b)** A representative FOV in DS5 (lung sarcoid) showing granulomatous lesion in lung sarcoidosis. Left to right, H&E image, patch-level saliency map (blue and orange, as indicated), inset, token-level saliency (1) for epithelioid histiocyte and (2) MNGC regions.

**Supplementary Figure 12. Resolving disagreement between human and SAE annotation**

**(a)** Representative H&E WSI image in DS1 annotated by SME (lymphoid aggregate, green) and SAE (F59985, red). Insets, SAE-only (FP1, FP2) and SME-only (FN1, FN2) are indicated (arrows, boxes). **(b)** Top (left to right), H&E inset images FP1, FP2, FN1 and FN2 with annotations outlined (green, red). Bottom, corresponding token-level saliency (consolidated dictionary), as indicated.

