## Supplementary Information for "Compositional and interpretable representation of histology using AI foundation models and sparse autoencoders"

### Content


| Title | Page |
|---|---|
| Supplementary Figure 1: Defining histomorphologies by clustering FM patch embeddings | 3 |
| Supplementary Figure 2: Workflow for model training and human interpretation | 5 |
| Supplementary Figure 3: SAE model training and evaluation | 7 |
| Supplementary Figure 4: SAE feature grouping and consolidated token labels in DS1 | 9 |
| Supplementary Figure 5: SAE features deconvolve tissue regions with overlapping histomorphologies | 11 |
| Supplementary Figure 6: Consistent feature representation in multiple sections of the same tissue block | 12 |
| Supplementary Figure 7: Alternative SAE algorithm provides broadly similar features | 13 |
| Supplementary Figure 8: Diversity of MNGCs in H&E images | 14 |
| Supplementary Figure 9: Accurate feature recall in multiple datasets | 16 |
| Supplementary Figure 10: Extended results demonstrating the generalizability of SAE features in other TB specimens | 17 |
| Supplementary Figure 11: Extended results demonstrating the generalizability of SAE features in other lung disease specimens | 18 |
| Supplementary Figure 12: Resolving disagreement between human and SAE annotation | 19 |
| Supplementary Table 1: Dataset metadata for DS1-5 | 21 |
| Supplementary Table 2: Interpretations for histomorphologic features defined from the K-means (K=25) clusters | 22 |
| Supplementary Table 3: Mapping between SAE1 feature index and feature groups | Separate Excel file |
| Supplementary Table 4: Interpretations for histomorphologic features defined from SAE1 feature groups | 24 |
| Supplementary Note | 25 |

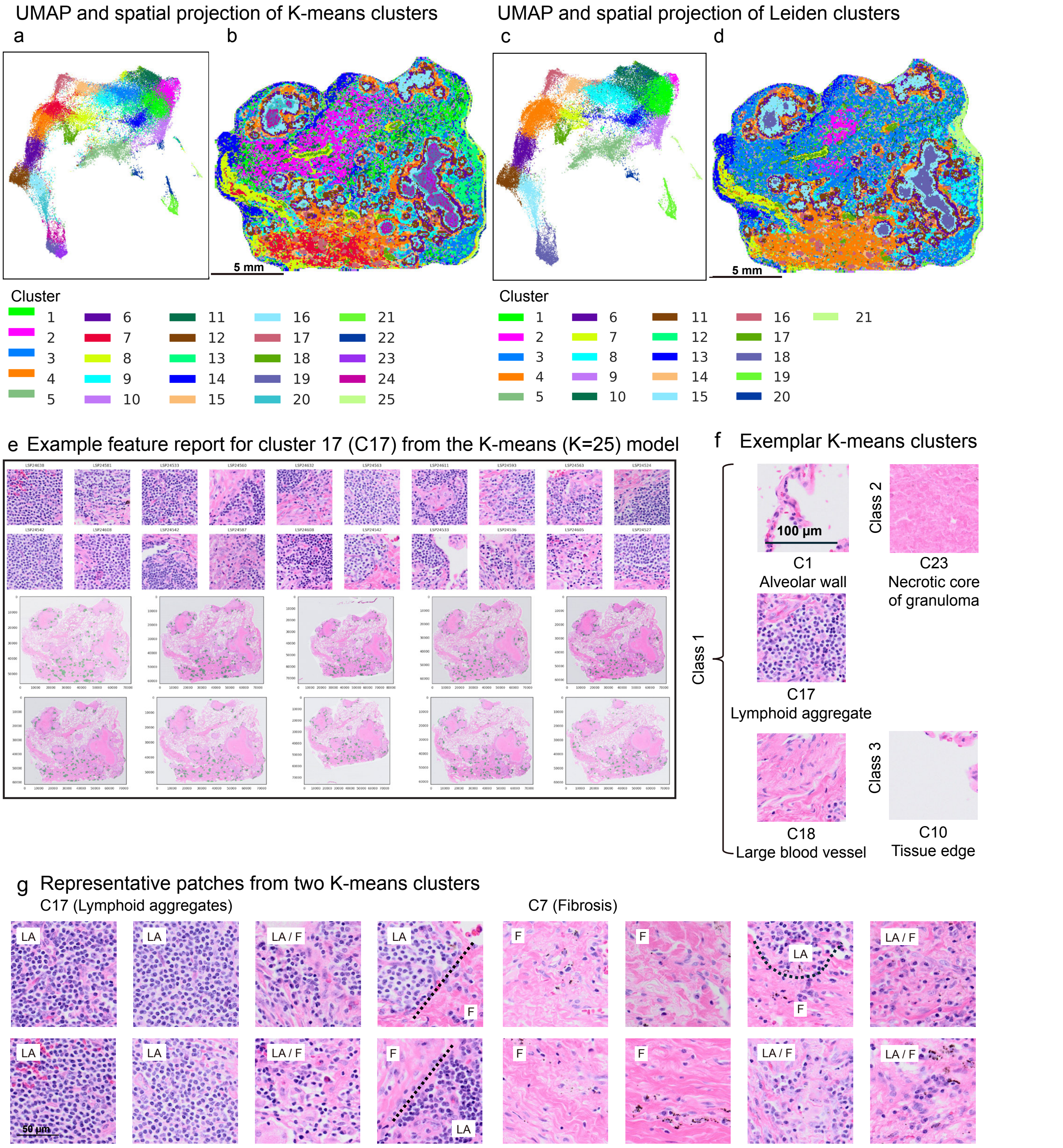


**Supplementary Figure 1. Defining histomorphologies by clustering FM patch embeddings**

**(a)** UMAP, sub-sampled (n=50,000) patch embeddings from UNI FM. Color represents patch assignment to K-means clusters (K=25). **(b)** Multi-feature maps for a representative tissue section using K-means with colors as in **(a)**. **(c)** UMAP for the same sub-sampled patch set. Color represents patch assignment to Leiden clusters (n=21). **(d)** Multi-feature maps the same section using Leiden algorithm with colors as in **(c)**. **(e)** Feature report for cluster C17. Top, 20 representative H&E image patches with source WSI indicated, bottom, 10 WSI H&E feature maps with x and y pixel positions indicated. **(f)** Representative image patches for K-means clusters classified based on morphology as class 1: cellular structure, class 2: homogenous, acellular material, class 3: blank space and debris. Right, cluster indices and interpretations shown. **(g)** Selected representative H&E image patches from C17 (lymphoid aggregate, LA) and C7 (fibrosis, F) annotated for regions of F or LA, with boundary indicated (dashed). LA/F indicates FOV with no clear boundary.

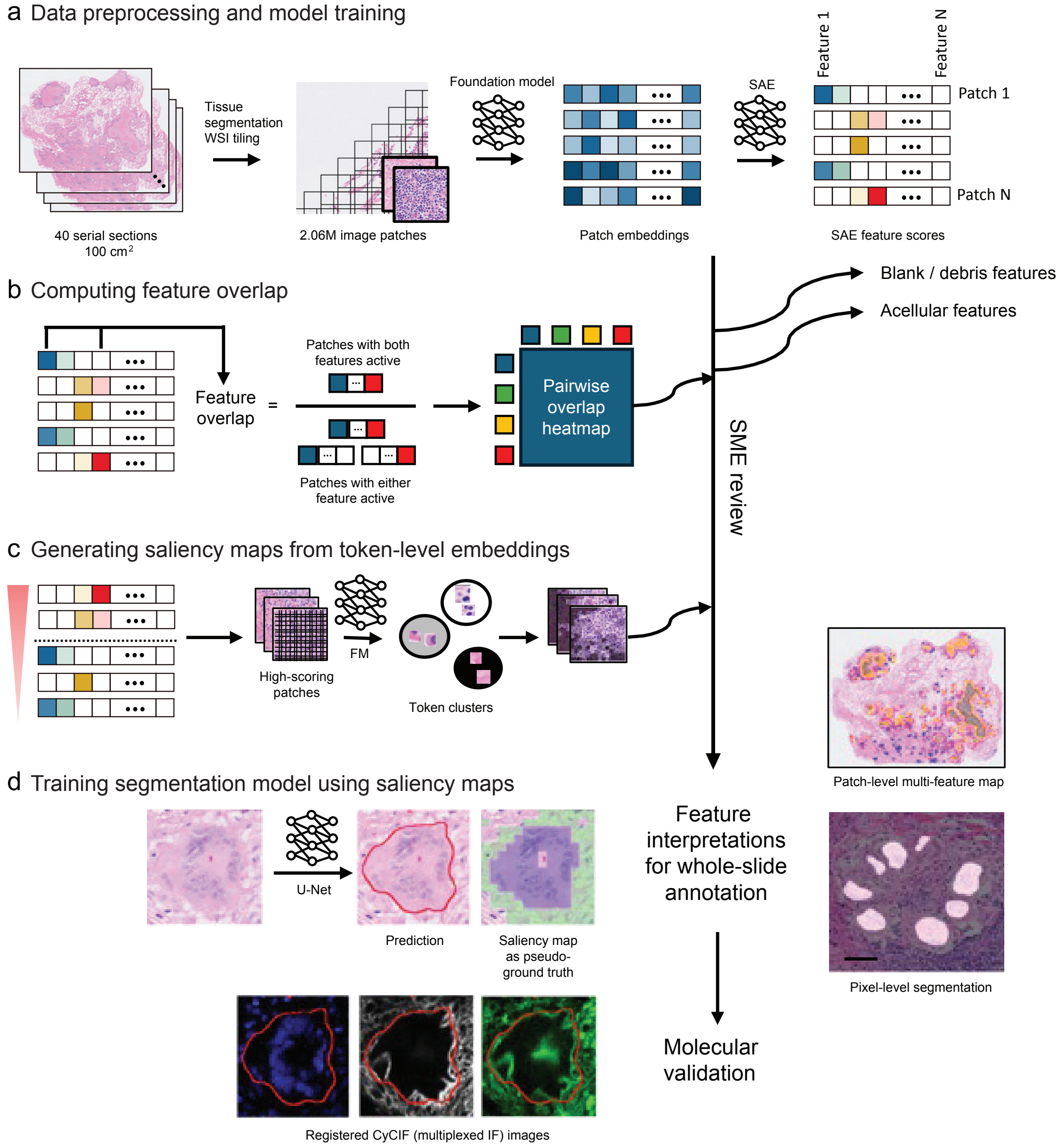


**Supplementary Figure 2. Workflow for model training and human interpretation**

**(a)** Initial dataset preprocessing and model training. H&E WSIs are tiled into overlapping patches and embedded using a FM (UNI). A one-layer SAE is trained to reconstruct the FM embeddings using sparse features. Only active features are kept (**Supplementary Fig. 3a**, methods). After training, all FM embeddings are encoded by the trained model into SAE features. Right: each row is the FM embedding of one image patch and each column is an SAE feature. For any patch, most features are not active because of the feature sparsity imposed during model training. **(b)** For each pair of SAE feature, we compute feature overlap as the fraction of patches with high score for either feature that is high for both. The features are ordered by hierarchically clustering the spatial overlap matrix. Combined with feature reports (**Supplementary Fig. 3d**), SMEs could rapidly filter out features detecting blank space, debris, or acellular morphology. **(c)** For each SAE feature, patches are ranked by feature score, and high scoring patches are again embedded by the FM to produce token-level embeddings. Token-level clusters are mapped back to the patches to generate saliency maps to be included in feature reports. Most features are interpreted by this multiscale approach and grouped together, leading to multi-feature maps for whole-slide annotation. **(d)** For a particular feature of interest, a U-Net is trained to produce pixel segmentation that refines the patch level feature score by predicting relevant regions indicated by saliency maps. The trained U-Net is then applied across the whole slide, and the segmentation is transferred to adjacent section subjected to spatial profiling, such as with CyCIF, to validate the molecular morphology of the detected feature.

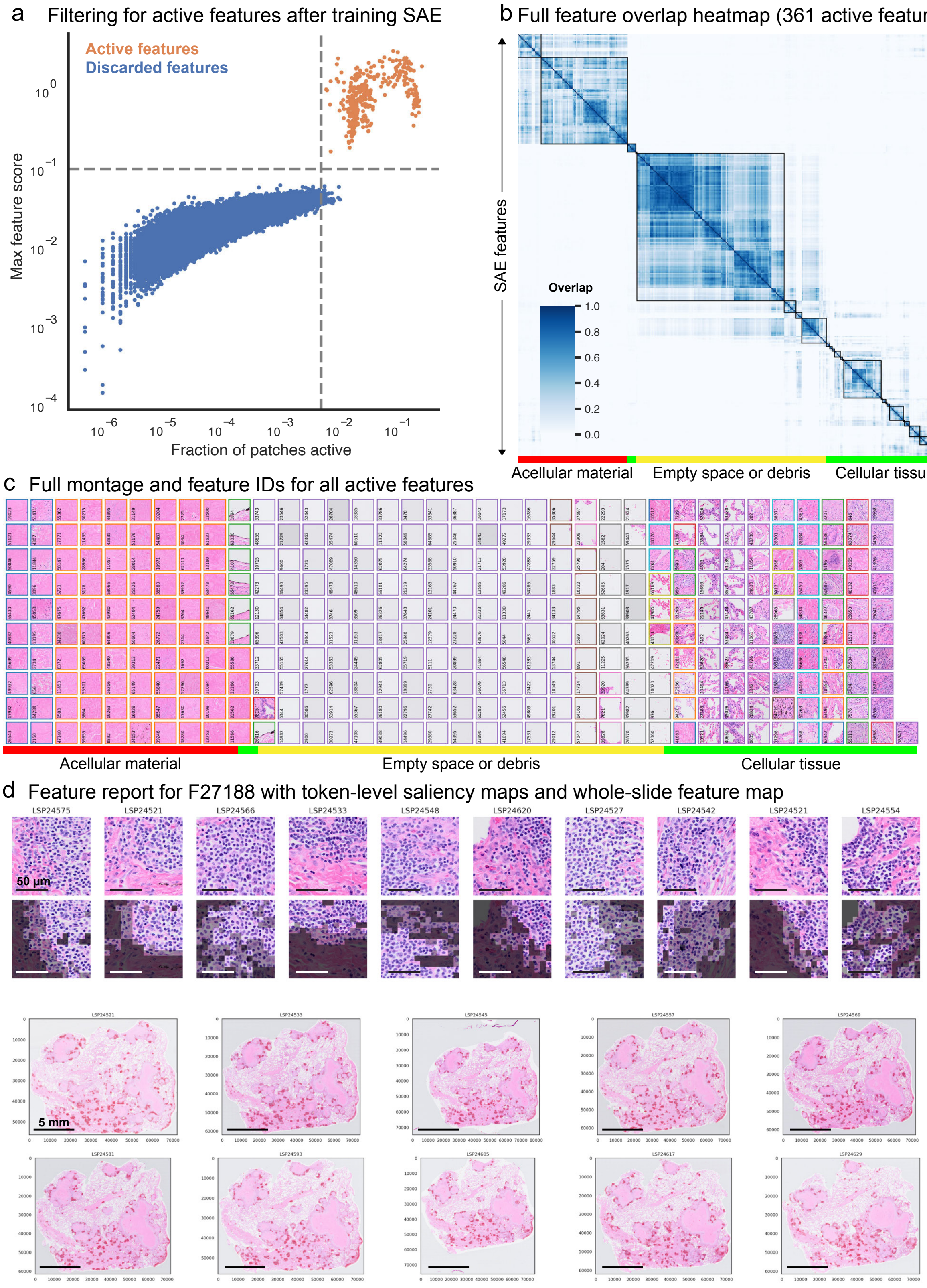


**Supplementary Figure 3. SAE model training and evaluation**

**(a)** Scatterplot, maximum feature score and fraction of active patches for all primary SAE features (n=65536). Features in the top right quadrant are considered active (orange). **(b)** Spatial overlap matrix for all 361 active features from SAE1 sorted by hierarchical clustering. Color bar (bottom) indicates visual classification of exemplar H&E image patches. **(c)** Montage of exemplar H&E image patches for all 361 active features from SAE1, as ordered in (b). Primary feature index (0-65535), right of each image. Color bar (bottom) indicates classification of features, as (b). **(d)** SAE feature report for F27188. Top, 10 highest scoring H&E image patches and, below, corresponding token-level saliency maps for K-means (K=3) clustered of FM embeddings. Bottom, 10 representative feature map, image patch scores projected by intensity (red). Source WSI for H&E image patches, feature maps, and x and y pixel coordinates indicated on axes of feature maps, as indicated.

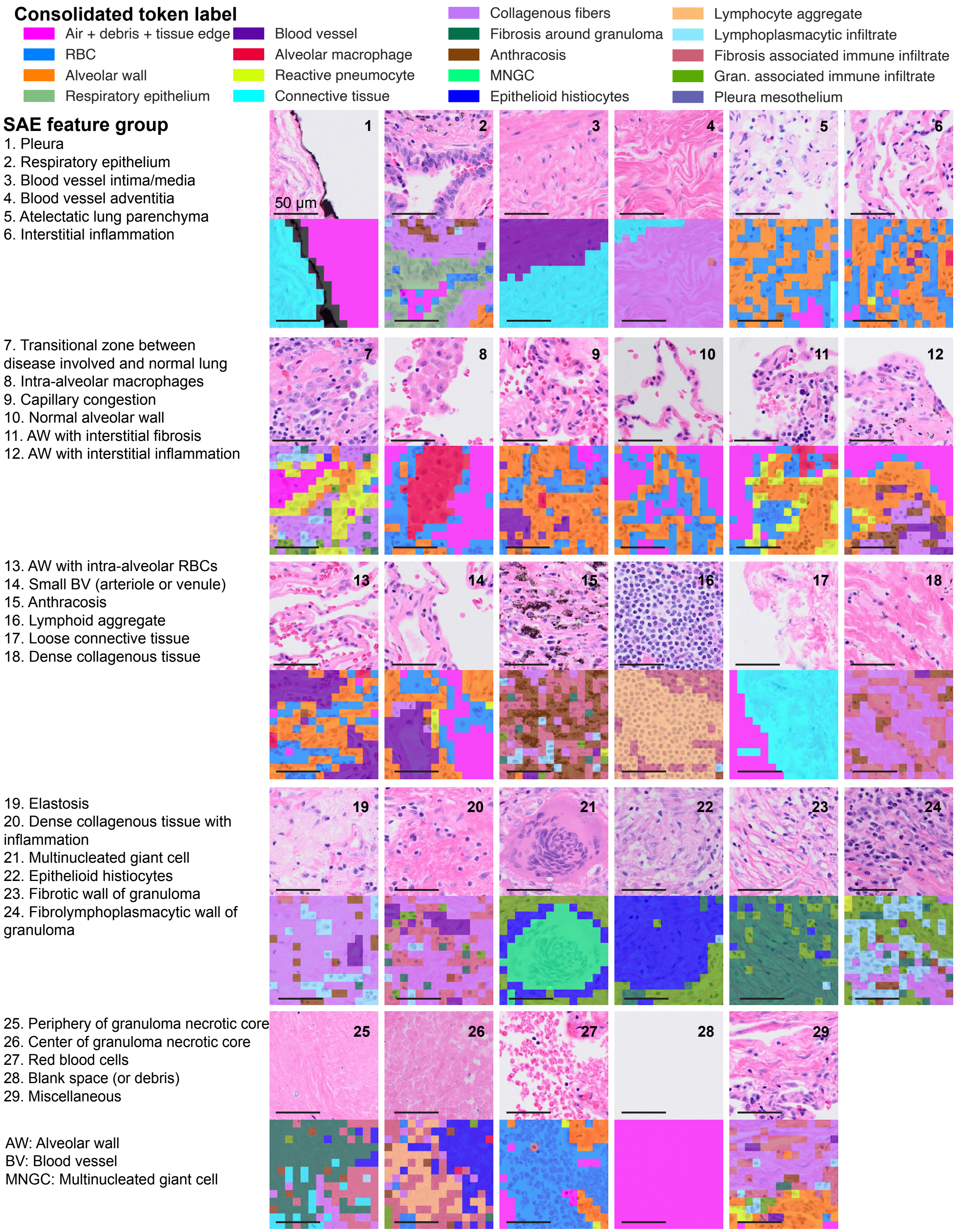


**Supplementary Figure 4. SAE feature grouping and consolidated token labels in DS1**

Exemplar H&E image patches (above) representing 29 groups of SAE features, and saliency maps (below) for token-level mapping of 18 distinct token groups. Left: feature group index, in order left-toright) and named histomorphology. Color coding and SME interpretation of token groups, as indicated (top).

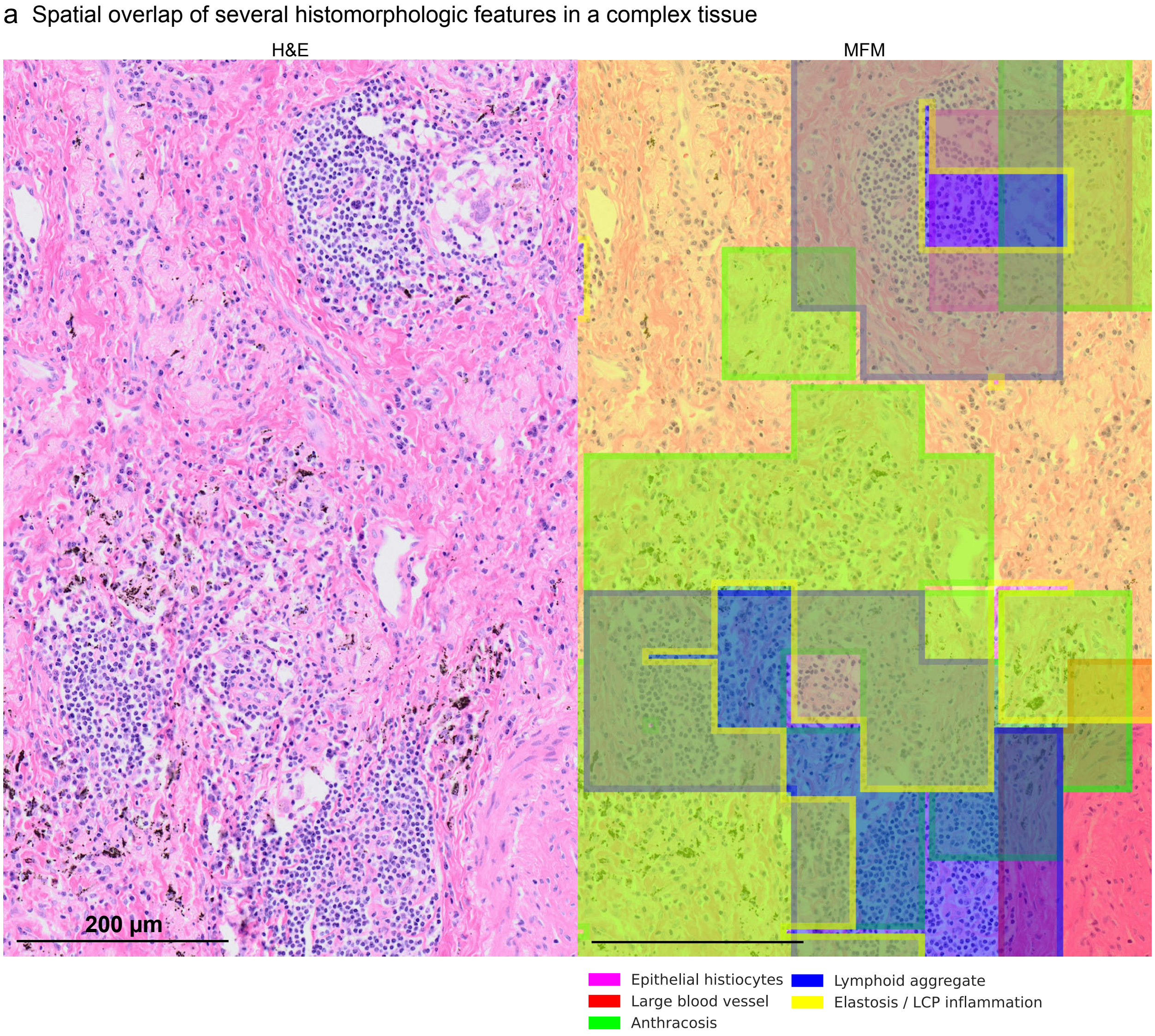


**Supplementary Figure 5. SAE features deconvolve tissue regions with overlapping histomorphologies**

H&E and corresponding MFM of FOV with multiple histomorphologies. Each feature map in MFM is binarized and has boundary colored solid to delineate regions that can overlap.

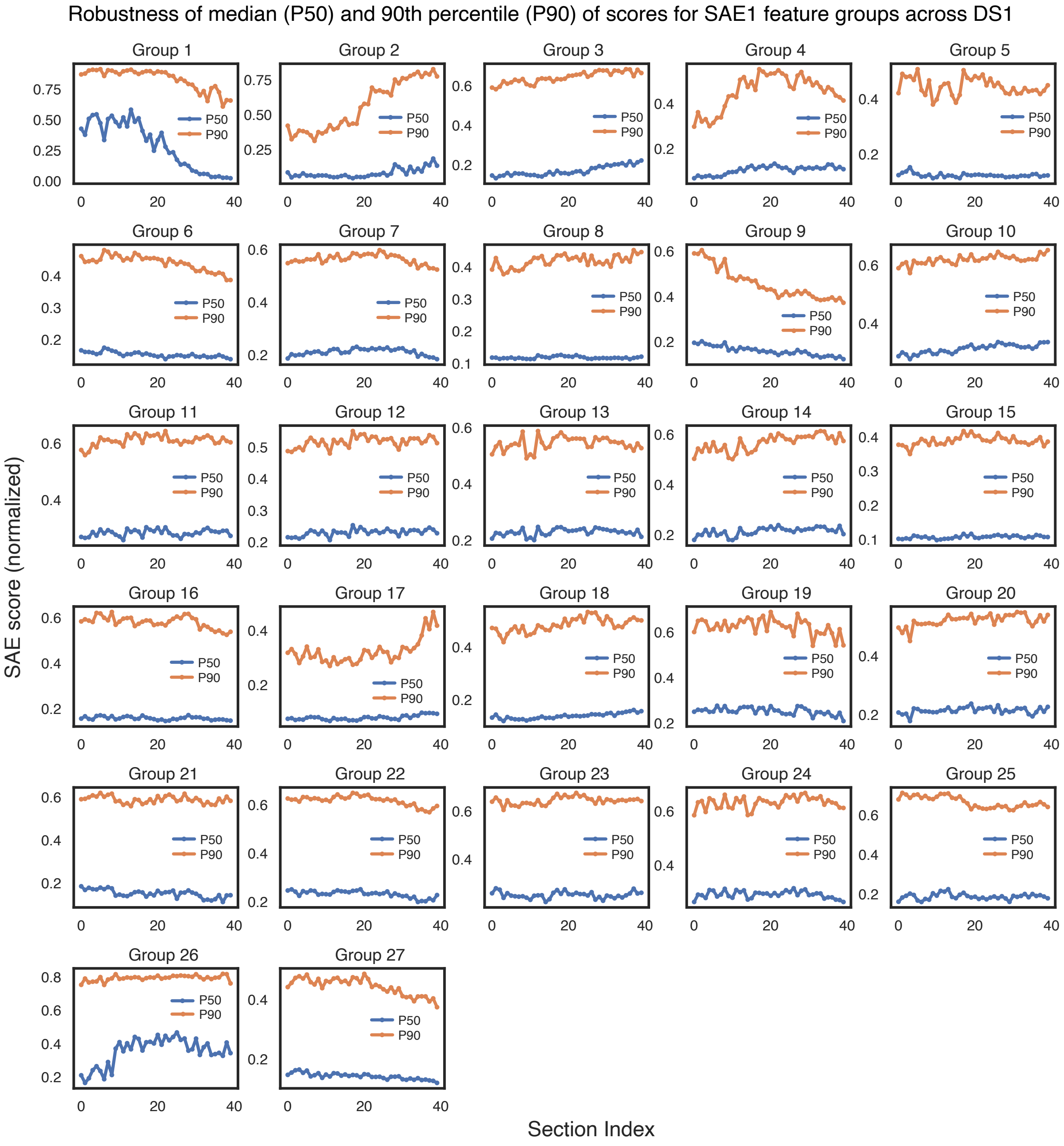


**Supplementary Figure 6. Consistent feature representation in multiple sections of the same tissue block**

For each SAE feature group, the median (blue) and 90$^{th}$ percentile (orange) of aggregated feature score is plotted across 40 consecutive sections from the tissue block.

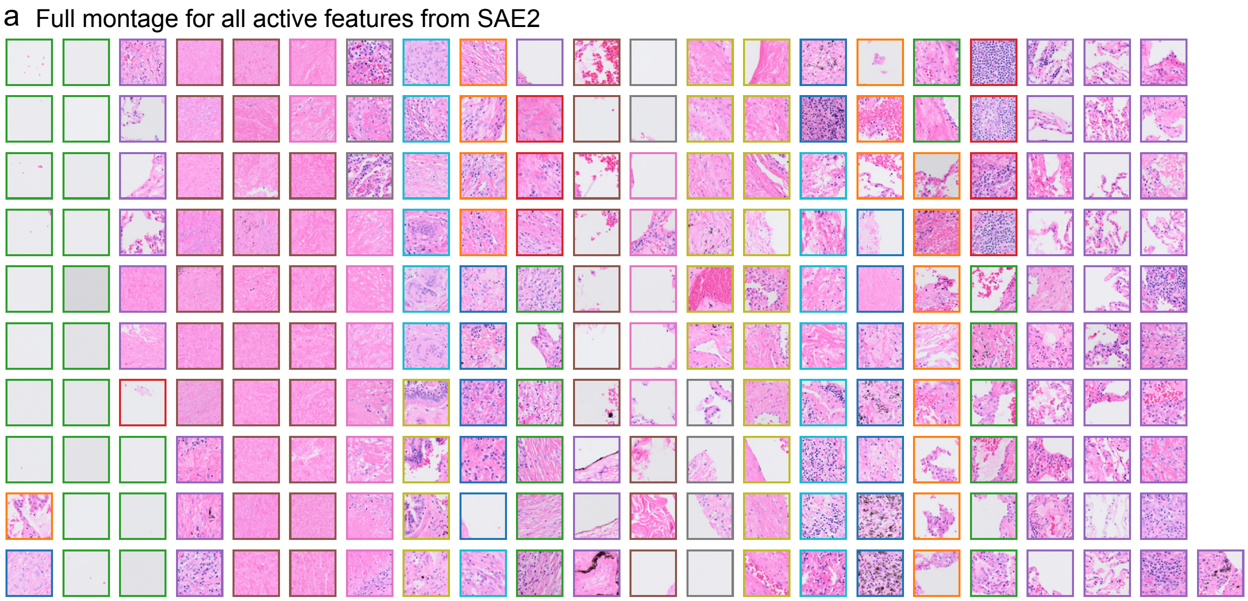


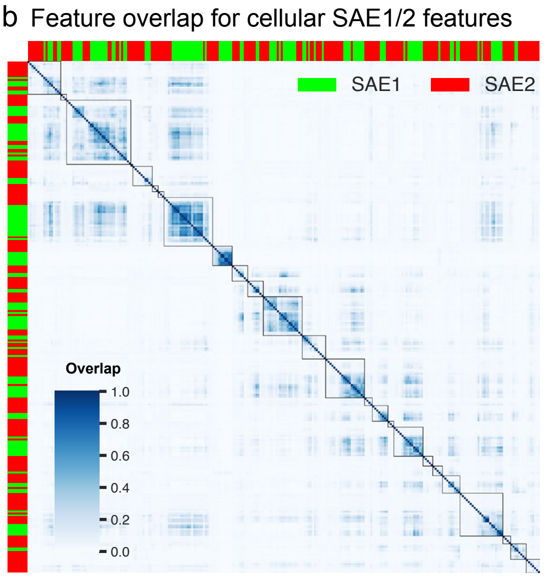


c Frequency of cellular features in each type of clusters

| **Cluster type** | SAE1 only | SAE2 only | SAE1 + SAE2 |
|---|---|---|---|
| **Frequency** | 0 (0%) | 33 (12%) | 230 (87%) |

**Supplementary Figure 7. Alternative SAE algorithm provides broadly similar features**

**(a)** Montage, H&E image patch exemplars of all SAE2 features. Image border color coded by clustering feature overlap similarly as in **Fig. 1c**. **(b)** Spatial overlap matrix of cellular SAE1 (n=110 out of 361) and SAE2 (n=153 out of 211) features grouped into 25 clusters by hierarchical clustering using complete linkage. Color bar (right or top) indicates feature from SAE1 (green) or SAE2 (red). **(c)** Frequency of cellular SAE1 and SAE2 features in each of the three types of clusters, depending on whether the cluster contain only SAE1 features, only SAE2 features, or a mixture of SAE1 and SAE2 features.

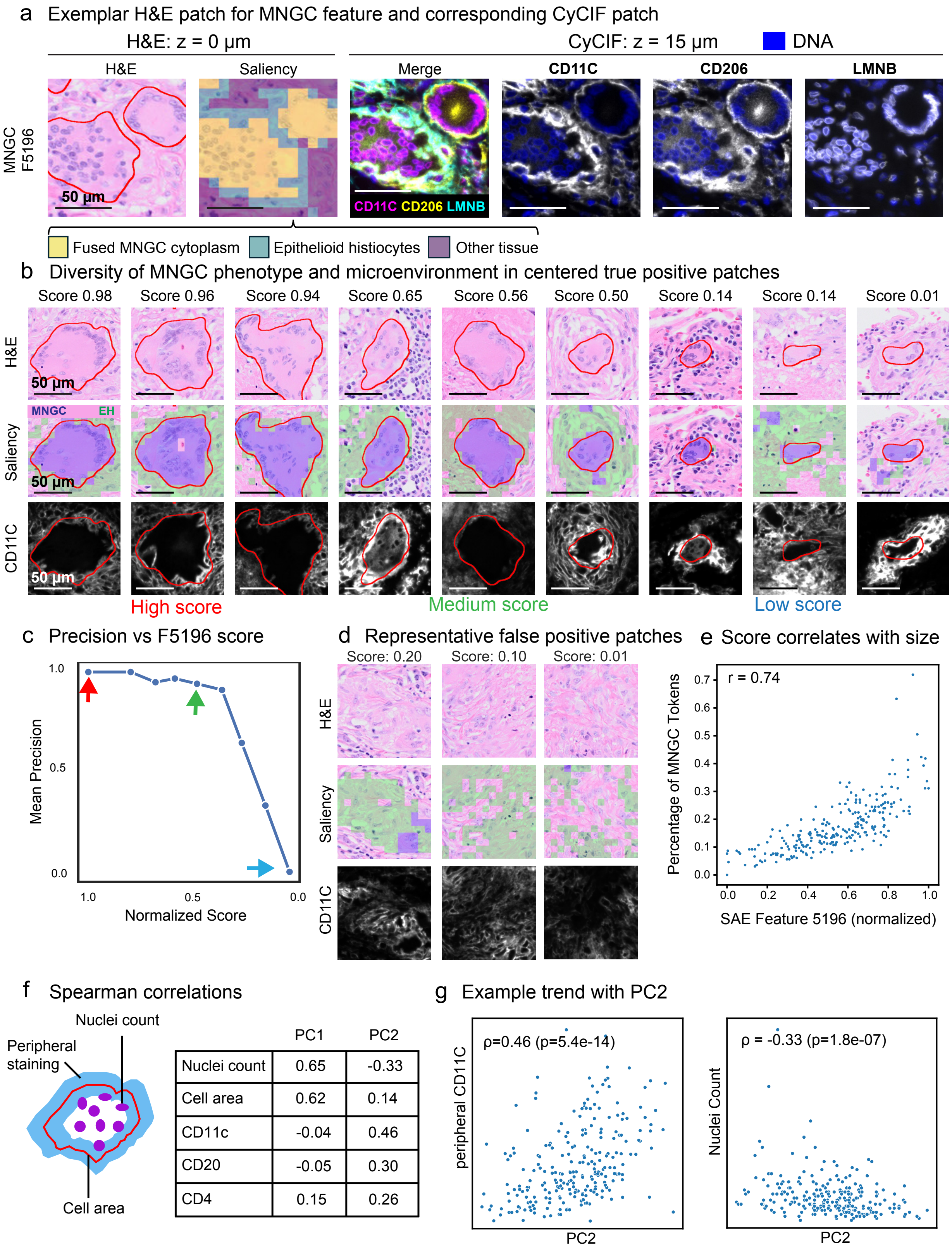


**Supplementary Figure 8. Diversity of MNGCs in H&E images**

**(a)** Exemplar H&E image patch for feature F5196 (MNGC). Left to right:  H&E image, feature-specific saliency map (K=3), multi-channel merge, and individual channels are shown on the same row. **(b)** Top row, three representative H&E patches selected of high, intermediate, and low F5196 feature score; middle: saliency map with consolidated dictionary, MNGC (violet) and epithelioid histiocytes (green); bottom: CD11c staining in registered, adjacent section (z-offset +5 μm). U-Net derived MNGC segmentation boundary is indicated (red) in all images. **(c)** Precision versus feature score for F5196. Arrows are color-coded to match (b) to illustrate top, medium, and low scoring patches. **(d)** Representative patches with low score for F5196 without MNGCs. The H&E patch, saliency map, and CD11c staining are shown in columns similarly to (b). **(e)** Scatterplot, correlation (Spearman: 0.74), of feature score and fraction of MNGC tokens. **(f)** Spearman correlation of PC1, PC2 and spatial features of U-Net segmented MNGCs in adjacent CyCIF image: cell area (red), nuclei within MNGC mask (purple) and mean fluorescent intensity for CD4, CD11C and CD20 at mask periphery (light blue). **(g)** Scatterplot, PC2 vs. CyCIF feature from (f) for MNGC-centered image patches. Spearman's rank correlation coefficient and p-value as indicated.

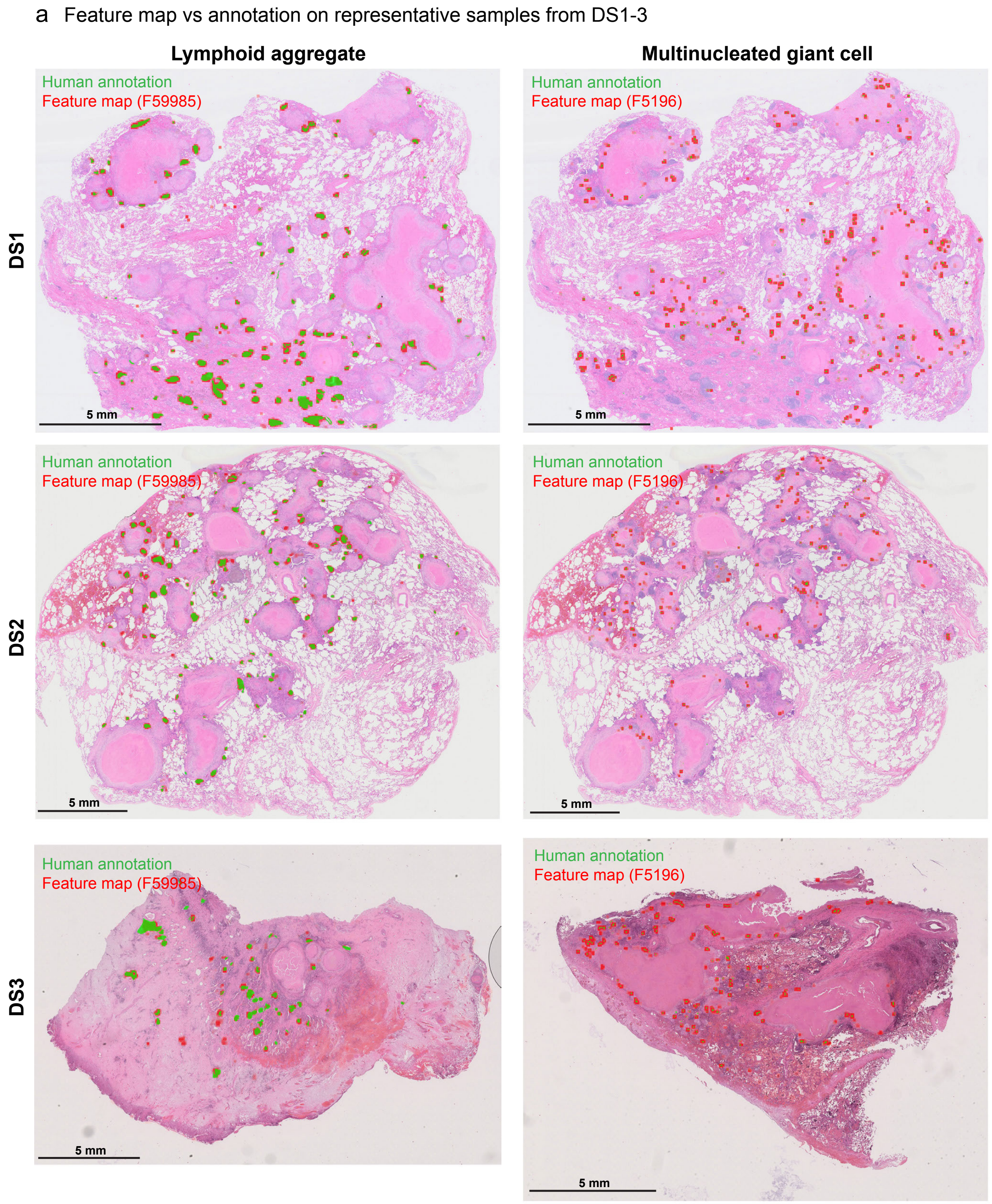


**Supplementary Figure 9. Accurate feature recall in multiple datasets**

For each combination of dataset from DS1 – 3 and histomorphologies from LA and MNGC, the whole-slide H&E image of the representative section is shown with areas annotated by SMEs in green and SAE feature map in red. SAE feature indices in figure legend.

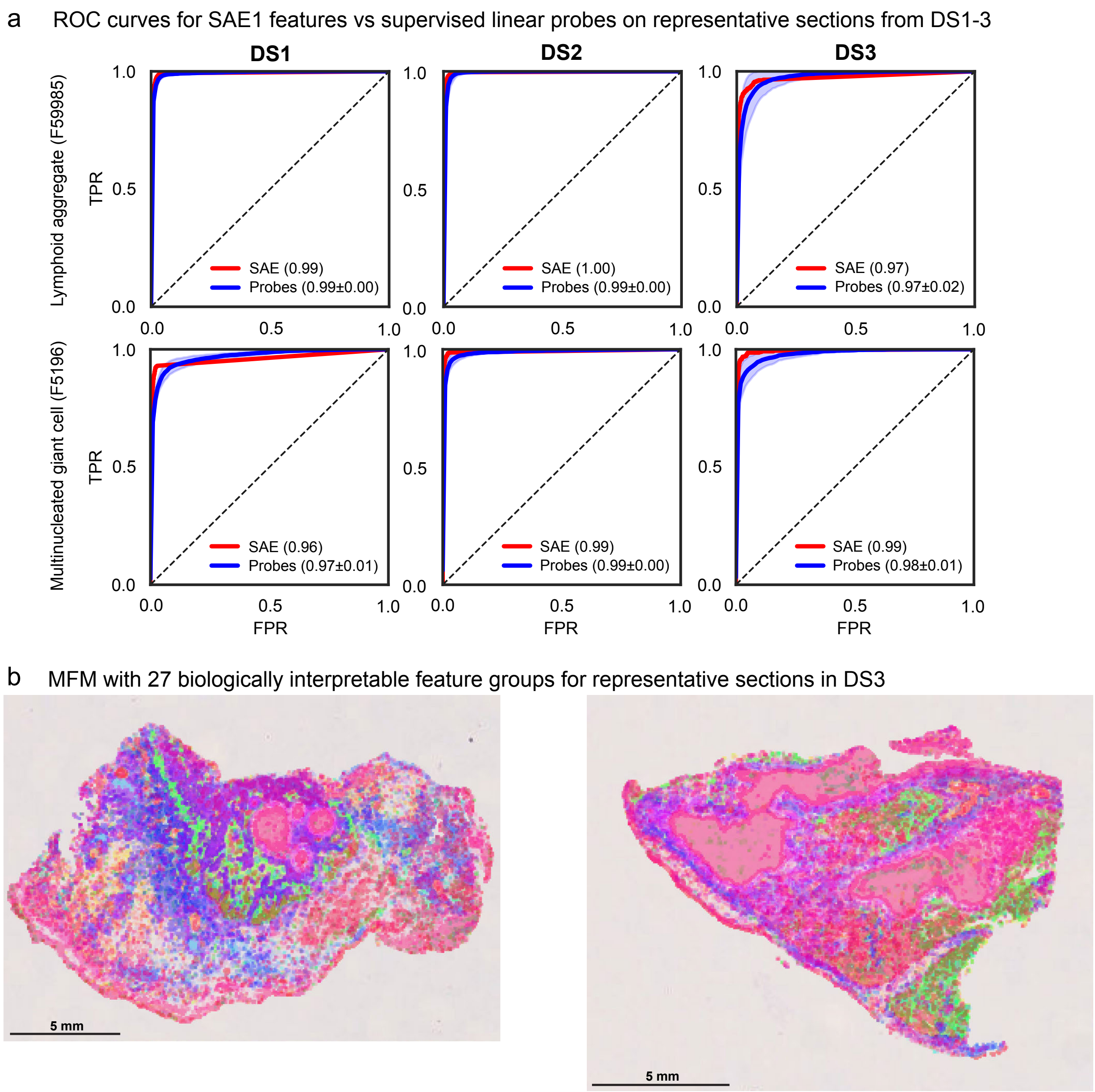


**Supplementary Figure 10. Extended results demonstrating the generalizability of SAE features in other TB specimens**

**(a)** Receiver operating characteristics (ROC) curves for SAE features (red) versus ensemble of linear probes (blue) for patch-level LA and MNGC classification in DS1 – 3. Average (solid line) and range (light zone). In parentheses, area under ROC curve (AUROC) for SAE and ensemble (average ± std. dev.). **(b)** MFMs, two representative H&E WSIs from DS4 projected score of 27 biologically interpretable groups of SAE1 features (same as **Fig. 3a**).

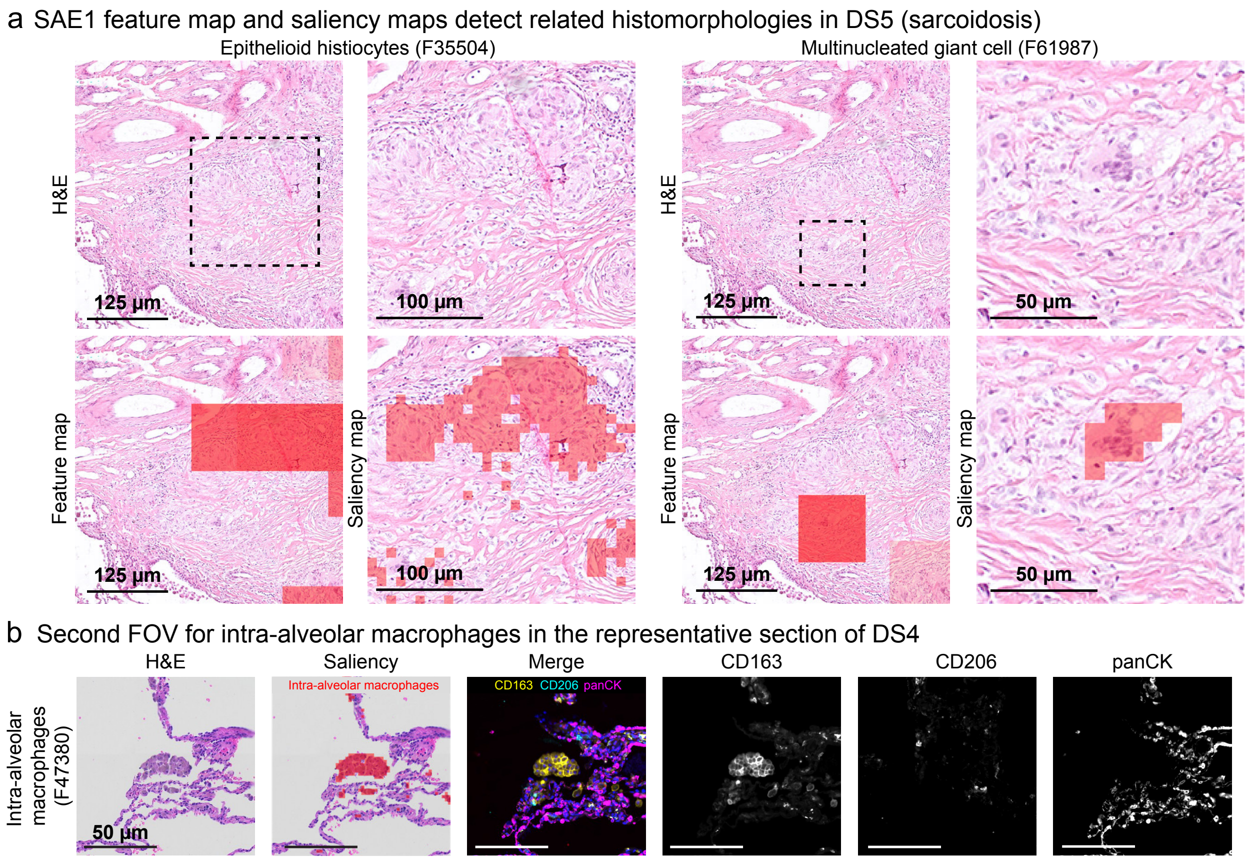


**Supplementary Figure 11. Extended results demonstrating the generalizability of SAE features in other lung disease specimens**

**(a)** FOV, alveolar macrophage (F47380) in DS4 (LUAD) similar to DS1. Left to right, H&E image patch, token level saliency map (red, consolidated dictionary), multi-channel merge, and single-channel grayscale images. **(b)** A representative FOV in DS5 (lung sarcoid) showing granulomatous lesion in lung sarcoidosis. Left to right, H&E image, patch-level saliency map (blue and orange, as indicated), inset, token-level saliency (1) for epithelioid histiocyte and (2) MNGC regions.

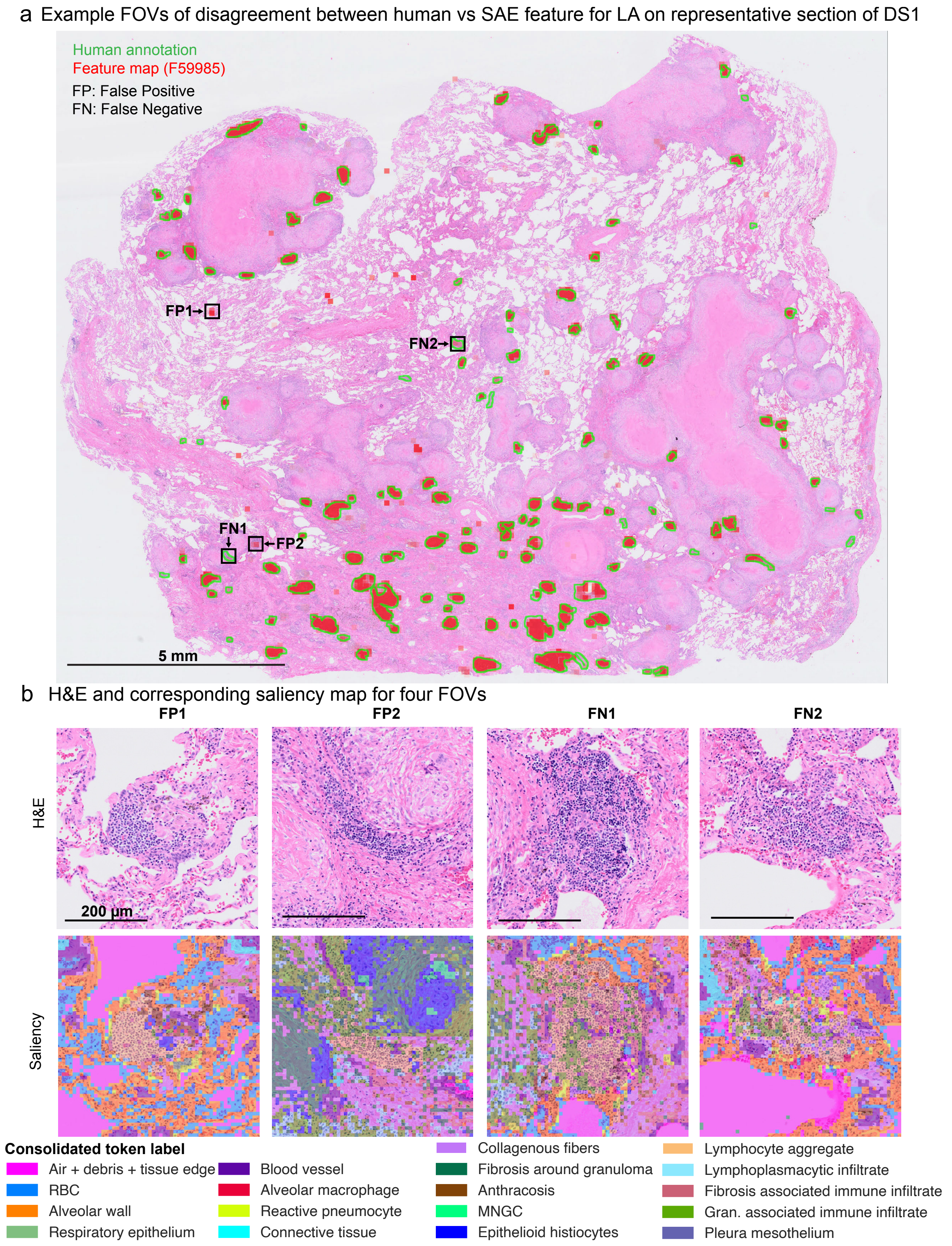


**Supplementary Figure 12. Resolving disagreement between human and SAE annotation**

**(a)** Representative H&E WSI image in DS1 annotated by SME (lymphoid aggregate, green) and SAE (F59985, red). Insets, SAE-only (FP1, FP2) and SME-only (FN1, FN2) are indicated (arrows, boxes).

**(b)** Top (left to right), H&E inset images FP1, FP2, FN1 and FN2 with annotations outlined (green, red). Bottom, corresponding token-level saliency (consolidated dictionary), as indicated.

| Dataset name | Disease type | Number of H&E WSIs | Medical center |
|---|---|---|---|
| DS1 | Pulmonary TB | 40 | Brigham and Women's Hospital |
| DS2 | Pulmonary TB | 15 | Brigham and Women's Hospital |
| DS3 | Pulmonary TB | 20 | University of KwaZulu-Natal (AHRI) |
| DS4 | Lung adenocarcinoma | 12 | Brigham and Women's Hospital |
| DS5 | Sarcoidosis | 5 | Stockholm University and Research Center Borstel |

**Supplementary Table 1**: Dataset metadata for DS1-5 used in the current study.

| K-means cluster ID | Class | Tissue compartment | Attempted interpretation |
|---|---|---|---|
| 0 | Cellular | Alveoli | Transitional zone |
| 1 | Cellular | Alveoli | Alveolar interstitium |
| 2 | Cellular | Connective tissue | Mixed connective tissue |
| 3 | Cellular | Connective tissue | Transitional zone |
| 4 | Cellular | Undefined | Undefined |
| 5 | Cellular | Undefined | Undefined |
| 6 | Cellular | Granuloma | Lymphocytic cuff |
| 7 | Cellular | Granuloma | Transitional zone |
| 8 | Cellular | Alveoli | Alveolar airspace |
| 9 | Cellular | Alveoli | Alveolar interstitium |
| 10 | Cellular | Alveoli | Alveolar interstitium |
| 11 | Cellular | Granuloma | Fibrotic cuff |
| 12 | Cellular | Alveoli | Alveolar interstitium with anthracosis |
| 13 | Cellular | Granuloma | Necrotic margin |
| 14 | Cellular | Granuloma | Myeloid cuff |
| 15 | Cellular | Connective tissue | Fibrosis with Anthracosis |
| 16 | Cellular | Lymphoid | Lymphoid aggregate |
| 17 | Cellular | Myeloid | Multinucleated giant cells and cellular granulomas |
| 18 | Cellular | Connective tissue | Large blood vessels and pleura |
| 19 | Cellular | Granuloma | Necrotic margin |
| 20 | No tissue | No tissue | No tissue |
| 21 | Acellular | Granuloma | Necrotic core |
| 22 | Acellular | Granuloma | Necrotic core |
| 23 | No tissue | No tissue | No tissue |
| 24 | Acellular | Granuloma | Necrotic core |

**Supplementary Table 2**: Interpretations for histomorphologic features defined from the K-means (K=25) clusters.

**Supplementary Table 3**: Mapping between SAE1 feature index (1-65536) and feature groups (1-29).

| SAE feature group ID | Interpretation | # of features |
|---|---|---|
| 1 | Pleura | 7 |
| 2 | Respiratory epithelium | 1 |
| 3 | Blood vessel intima/media | 2 |
| 4 | Blood vessel adventitia | 2 |
| 5 | Atelectatic lung parenchyma | 2 |
| 6 | Alveolar wall with interstitial inflammation | 2 |
| 7 | Transitional zone between disease involved and normal lung | 2 |
| 8 | Intra-alveolar macrophages | 1 |
| 9 | Capillary congestion | 1 |
| 10 | Normal alveolar wall | 6 |
| 11 | Alveolar wall with interstitial fibrosis | 16 |
| 12 | Alveolar wall with interstitial inflammation | 4 |
| 13 | Alveolar wall with intra-alveolar RBCs | 1 |
| 14 | Small BV (arteriole or venule) | 4 |
| 15 | Anthracosis | 1 |
| 16 | Lymphoid aggregate | 3 |
| 17 | Loose connective tissue | 1 |
| 18 | Dense collagenous tissue | 2 |
| 19 | Elastosis | 5 |
| 20 | Dense collagenous tissue with inflammation | 7 |
| 21 | Multinucleated giant cell | 4 |
| 22 | Epithelioid histiocytes | 5 |
| 23 | Fibrotic wall of granuloma | 7 |
| 24 | Fibrolymphoplasmacytic wall of granuloma | 10 |
| 25 | Periphery of granuloma necrotic core | 20 |
| 26 | Center of granuloma necrotic core | 74 |
| 27 | RBCs | 4 |
| 28 | Blank space (or debris) | 162 |
| 29 | Miscellaneous | 5 |
| | **Total** | **361** |

**Supplementary Table 4**: Interpretations for histomorphologic features defined from SAE1 feature groups.

#### SUPPLEMENTARY NOTE

##### Considerations in human-machine collaboration in analysis of histology images

A key feature of this manuscript is an effort to align human knowledge of histopathology with classifications made by an FM-SAE computational model. Since our goal is ultimately to develop a practical approach to AI-enabled tissue annotation and analysis, our work raises the question: in what way does model-based analysis improve on, or complement, existing human approaches. Addressing this question rigorously will require formal software evaluation[1], likely coupled to an inter-rater reliability study[2], both of which are substantial efforts that we have not yet undertaken (see the limitations section of the discussion). However, because our work involved multiple pathologists who interacted frequently with each other and with the computational team, it is possible to make some preliminary observations.

The most obvious advantage of a reliable computational tool is that it is rapid and low-cost. Whole-slide histopathology and spatial profiling studies commonly involve dozens to hundreds of specimens, and thorough annotation of even a single recurrent feature (such as lymphoid aggregate or giant cell) is an onerous task.

Consistency is a second obvious advantage of a computational method: much of the disagreement between SMEs involves edge cases in which the correct call is not obvious (**Supplementary Fig. 12**). In these cases, model-based approaches may not be more accurate than humans, but should be more consistent. As additional data become available (single-cell spatial profiles for example), it will be possible to leverage consistency in annotation to better understand transitional histological states[3,4]. Computational tools are also expected to help reduce inter-observer variability in diagnostic tasks[4]. A key step in realizing these benefits will be constructing curated feature dictionaries; the dictionaries in this study represent only an initial effort in this direction.

A third advantage of computational modeling is that it can be applied in settings in which histopathology resources are limited. This situation prevails in many molecular spatial profiling studies (“spatial proteomics”)[5] which remain remarkably disconnected from traditional histopathology (outside of efforts to predict molecular features from H&E images[6]). This occurs because pathologists do not have sufficient research time available for detailed review of large numbers of slides. The LungInsight Collaboration[7] has identified accurate histological analysis as a substantial challenge in preclinical respiratory disease models. The availability of reliable AI tools for H&E annotation and analysis workflow should make this critical aspect of spatial biology more widely availability.

Our group also concluded the even domain experts – thoracic pathologists – benefitted from the ability of FM-SAE models to detect subtle correlations between spatial features, as well as morphological variation within a single complex feature. We illustrate this for MNGCs, which have a canonical appearance in only a subset of images but can be difficult to detect in others. Further study of this issue should make it possible to determine how practitioners can best utilize automated tools during routine diagnostic evaluation.
